# Generalizability of “GWAS hits” in clinical populations: Lessons from childhood cancer survivors

**DOI:** 10.1101/2020.02.02.930818

**Authors:** Cindy Im, Na Qin, Zhaoming Wang, Weiyu Qiu, Carrie R. Howell, Yadav Sapkota, Wonjong Moon, Wassim Chemaitilly, Todd M. Gibson, Daniel A. Mulrooney, Kirsten K. Ness, Carmen L. Wilson, Lindsay M. Morton, Gregory T. Armstrong, Smita Bhatia, Jinghui Zhang, Melissa M. Hudson, Leslie L. Robison, Yutaka Yasui

## Abstract

With mounting interest in translating GWAS hits from large meta-analyses (meta-GWAS) in diverse clinical settings, evaluating their generalizability in target populations is crucial. Here we consider long-term survivors of childhood cancers from the St. Jude Lifetime Cohort Study and show the limited generalizability of 1,376 robust SNP associations reported in the general population across 12 complex anthropometric and cardiometabolic phenotypes (N=2,231; observed-to-expected replication ratio=0.68, *P*=2.4×10^−9^). An examination of five comparable phenotypes in a second independent cohort of survivors from the Childhood Cancer Survivor Study corroborated the overall limited generalizability of meta-GWAS hits to survivors (N=4,212, observed-to-expected replication ratio=0.53, *P*=1.1×10^−16^). Meta-GWAS hits were less likely to be replicated in survivors exposed to cancer therapies associated with phenotype risk. Examination of complementary DNA methylation data in a subset of survivors revealed that treatment-related methylation patterns at genomic sites linked to meta-GWAS hits may disrupt established genetic signals in survivors.

---

Recent meta-analyses of genome-wide association studies (meta-GWAS) with large study samples (N>10,000) have discovered novel and replicated known associations between common genetic variants (i.e., single nucleotide polymorphisms or SNPs) and many complex traits and diseases. Genetic associations reported in cohorts with individuals of predominantly European ancestry have proven to be highly generalizable in other European cohorts^1^. For example, a recent examination of genome-wide significant associations for 32 complex traits across five broad disease groups reported a median replication rate of 84% in a cohort with >13,000 individuals of European ancestry^2^.

The generalizability of robust genetic associations reported by large-scale meta-GWAS (hereafter referred to as meta-GWAS hits) from the general population to specialized clinical populations has not been established for most complex phenotypes. Yet there is growing enthusiasm for utilizing polygenic risk scores to predict disease risk and identify high-risk individuals for targeted interventions; for example, polygenic risk scores have been shown to improve clinical prediction models for cardiovascular disease risk and used to support pharmaceutical interventions to target reductions in low-density lipoprotein levels in high-risk individuals^1,3^. It is imperative to evaluate the generalizability of established meta-GWAS hits in target populations before adopting such genetic tools built on the GWAS literature. Childhood cancer survivors are one such example of a specialized clinical population that would greatly benefit from knowledge of the generalizability of meta-GWAS hits. Today, approximately one in every 750 individuals is a survivor of childhood or adolescent cancer in the United States^4^. This growing population of survivors differs markedly from the general population. Studies have consistently shown that survivors are at greater risk for a wide range of serious health conditions earlier in life relative to general population or sibling controls, in part due to their exposures to treatments necessary to cure pediatric cancers^4–8^, including chronic cardiovascular and metabolic health conditions that are among the leading causes of morbidity and mortality among survivors^5,9–12^.

Here we report on the limited generalizability of 1,376 robust meta-GWAS hits (*P*<5×10^−8^) identified from the literature for 12 anthropometric and cardiometabolic phenotypes to adult survivors of childhood cancer from the St. Jude Lifetime Cohort Study^7^ (SJLIFE; N=2,231, European ancestry), a single-institution retrospective cohort study with longitudinal follow-up of survivors with clinically ascertained health outcomes. We also found limited generalizability of meta-GWAS hits in a second cohort of survivors for five phenotypes available for comparison from the Childhood Cancer Survivor Study (CCSS; N=4,212, European ancestry), a multi-center study with self-reported health conditions. Depletions of replicated meta-GWAS hits were exacerbated in survivor subgroups exposed to certain cancer treatments, particularly when treatments had larger contributions to phenotype variation. Lastly, we conducted ancillary analyses to explore the role of DNA methylation, an epigenetic alteration that is influenced by both inherited genetic variation and environmental factors^13^. Among the 236 survivors of SJLIFE with both germline methylome and genotype data, we found that cancer treatments, particularly radiation therapy, may obscure some robust meta-GWAS SNP associations in survivors.

## RESULTS

### Compiling robust meta-GWAS hits

The 12 phenotypes of interest included three anthropometric traits (height, body mass index [BMI], waist-to-hip ratio [WHR]); two blood pressure traits (systolic [SBP], diastolic [DBP]); four serum lipid traits (high-density lipoprotein levels [HDL], low-density lipoprotein levels [LDL], total cholesterol levels [TC], triglycerides [TG]); and three cardiometabolic disease outcomes (coronary artery disease [CAD], obesity, type 2 diabetes [T2D]). Using the NHGRI-EBI GWAS Catalog^14^, we identified 149 GWAS for these 12 phenotypes. After reviewing the literature against criteria for relevance, ancestry, and study suitability (see Methods), we compiled 1,415 genome-wide significant (*P*<5×10^−8^) SNP-phenotype associations from 46 selected GWAS featuring meta-analyses with replication studies that included >10,000 participants of predominantly European ancestry (Figure 1). We limited our analysis to the 1,376 SNP-phenotype associations (97.2%) that could be directly tested using 1,231 SNPs measured in SJLIFE that passed strict quality control.

**Figure 1:**
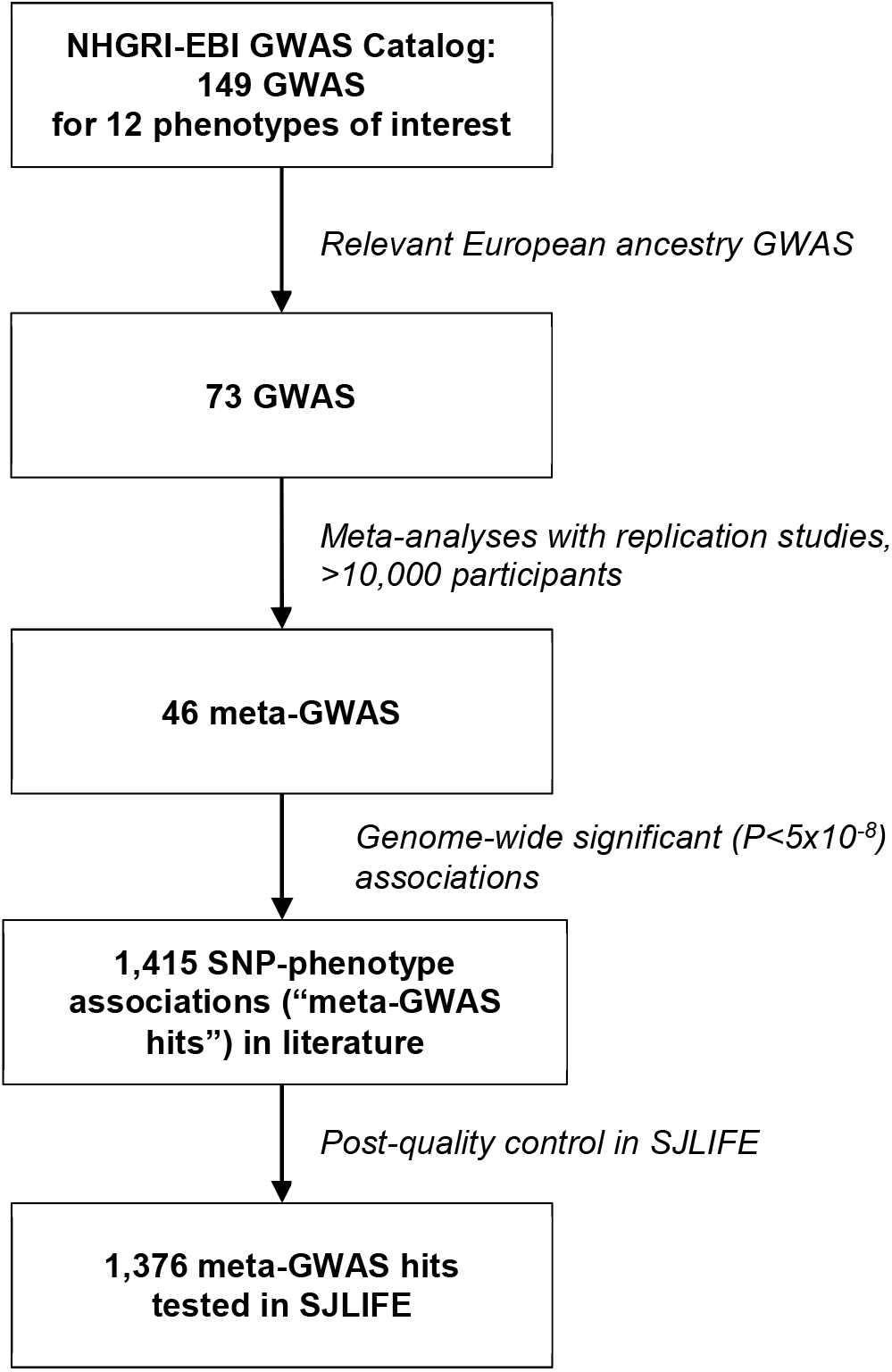
Diagram describing selection of meta-GWAS and genome-wide significant SNP-phenotype associations for replication in childhood cancer survivor cohorts. All reference GWAS considered in the current study were published between 1/1/2008 – 11/20/2017.

### Replicating meta-GWAS hits in SJLIFE childhood cancer survivors

Using phenotype definitions, adjustment covariates, and exclusion criteria that were consistent with reference GWAS (Table 1), our primary aim was to replicate the 1,376 robust meta-GWAS hits in 2,231 adult long-term (≥5-year) survivors of childhood cancer of European ancestry in SJLIFE^7^. Relevant descriptive statistics for the SJLIFE cohort are provided in Table 2. Most survivors had been exposed to at least one type of chemotherapeutic agent (85.3%) and over half (58.3%) had received radiotherapy; additional adjustments for specific cancer treatment exposures were considered based on the childhood cancer survivorship literature (Table 1). There was high correspondence between effect allele frequencies (EAFs) reported in the reference GWAS and the SJLIFE sample, with a median absolute difference of 0.99% (IQR=0.47-1.71%).

**Table 1:** Summary of methodological components for each SNP-phenotype association analysis in SJLIFE

| Phenotype | Phenotype transformation<sup>a</sup> | Unit or definition<sup>b</sup> | GWAS adjustment covariates<sup>a</sup> | Childhood cancer survivor adjustment covariates<sup>c</sup> | Exclusions<sup>b</sup> | Reference meta-GWAS (PMID) |
| --- | --- | --- | --- | --- | --- | --- |
| *Anthropometric* |  |  |  |  |  |  |
| Height | Sex-standardized Z-score | cm | Age, ancestry | Surgical procedures affecting spinal growth; scoliosis; hypothalamic-pituitary axis tumors; cranial or craniospinal radiation | Genetic syndromes, health conditions affecting stature<sup>d</sup> | 25282103, 20881960, 19570815, 19343178, 18952825, 18391952, 18391951, 18391950, 18193045 |
| Body mass index (BMI) | Inverse normal transformation of residuals | kg/m<sup>2</sup>; BMI adjusted for amputation | Age, age<sup>2</sup>, sex, ancestry | Hypothalamic-pituitary axis tumors; cranial radiation; glucocorticoids | None | 25673413, 24064335, 23669352, 22982992, 20935630, 19079261, 18454148 |
| Waist-to-hip ratio (WHR) | Inverse normal transformation of sex-standardized residuals | Ratio of waist and hip circumference (cm) | Age, age<sup>2</sup>, BMI, ancestry | Hypothalamic-pituitary axis tumors; cranial radiation; glucocorticoids | None | 28443625, 25673412, 20935629 |
| *Blood pressure* |  |  |  |  |  |  |
| Systolic blood pressure (SBP) | +15 mmHg with use of blood pressure lowering medications | mmHg | Age, age<sup>2</sup>, sex, BMI, ancestry | Abdominal, pelvic radiation | Prior myocardial infarction or heart failure | 28135244, 28739976, 26390057, 21909115, 19430483, 19430479 |
| Diastolic blood pressure (DBP) | +10 mmHg with use of blood pressure lowering medications | mmHg | Age, age<sup>2</sup>, sex, BMI, ancestry | Abdominal, pelvic radiation | Prior myocardial infarction or heart failure | Same as SBP |
| *Blood lipids* |  |  |  |  |  |  |
| High-density lipoprotein (HDL) | Inverse normal transformation of residuals | mg/dL | Age, age<sup>2</sup>, sex, ancestry | Hypothalamic-pituitary axis tumors; cranial radiation | Use of lipid-lowering medications | 24097068, 19060906 |
| Low-density lipoprotein (LDL) | Inverse normal transformation of residuals | mg/dL | Age, age<sup>2</sup>, sex, ancestry | Hypothalamic-pituitary axis tumors; cranial radiation | Use of lipid-lowering medications | Same as HDL |
| Total cholesterol (TC) | Inverse normal transformation of residuals | mg/dL | Age, age<sup>2</sup>, sex, ancestry | Hypothalamic-pituitary axis tumors; cranial radiation | Use of lipid-lowering medications | 24097068 |
| Triglycerides (TG) | Inverse normal transformation of residuals | mg/dL | Age, age<sup>2</sup>, sex, ancestry | Hypothalamic-pituitary axis tumors; cranial radiation | Use of lipid-lowering medications | Same as HDL |
| *Cardiometabolic disease* |  |  |  |  |  |  |
| Coronary artery disease (CAD) | None | Cases: CTCAE grades ≥2 | Age, sex, ancestry | BMI; smoking; cardiac-directed radiation; anthracyclines; platinum (cisplatin, carboplatin) | None | 28714975, 26950853, 26343387, 19198609 |
| Type 2 diabetes (T2D) | None | Cases: CTCAE grades ≥2 | Age, sex, BMI, ancestry | Cranial radiation; abdominal radiation | None | 28869590, 28566273, 24509480, 20581827, 20418489, 19734900, 18372903 |
| Obesity | None | Cases: BMI ≥30 kg/m<sup>2</sup> | Age, sex, ancestry | Hypothalamic-pituitary axis tumors; cranial radiation; glucocorticoids | None | 23563607, 21708048 |

Abbreviations: genome-wide association study (GWAS); cm (centimeter); kg (kilogram); m (meter); mmHg (millimeter of mercury); CTCAE (Common Terminology Criteria for Adverse Events, modified v4.03).
a. GWAS covariates, as defined by reference GWAS.
b. Phenotype units and definitions and participant exclusion criteria from reference GWAS were reviewed and adapted when necessary for analysis in SJLIFE.
c. Covariates specific to childhood cancer survivors, based on the childhood cancer survivorship research literature.
d. Syndromes and health conditions affecting height include: Down syndrome; Turner syndrome; Neurofibromatosis, type 1; Russell-Silver syndrome; benign bone lesion/cysts; cartilage disorder, skeletal spine disorder.

**Table 2:**
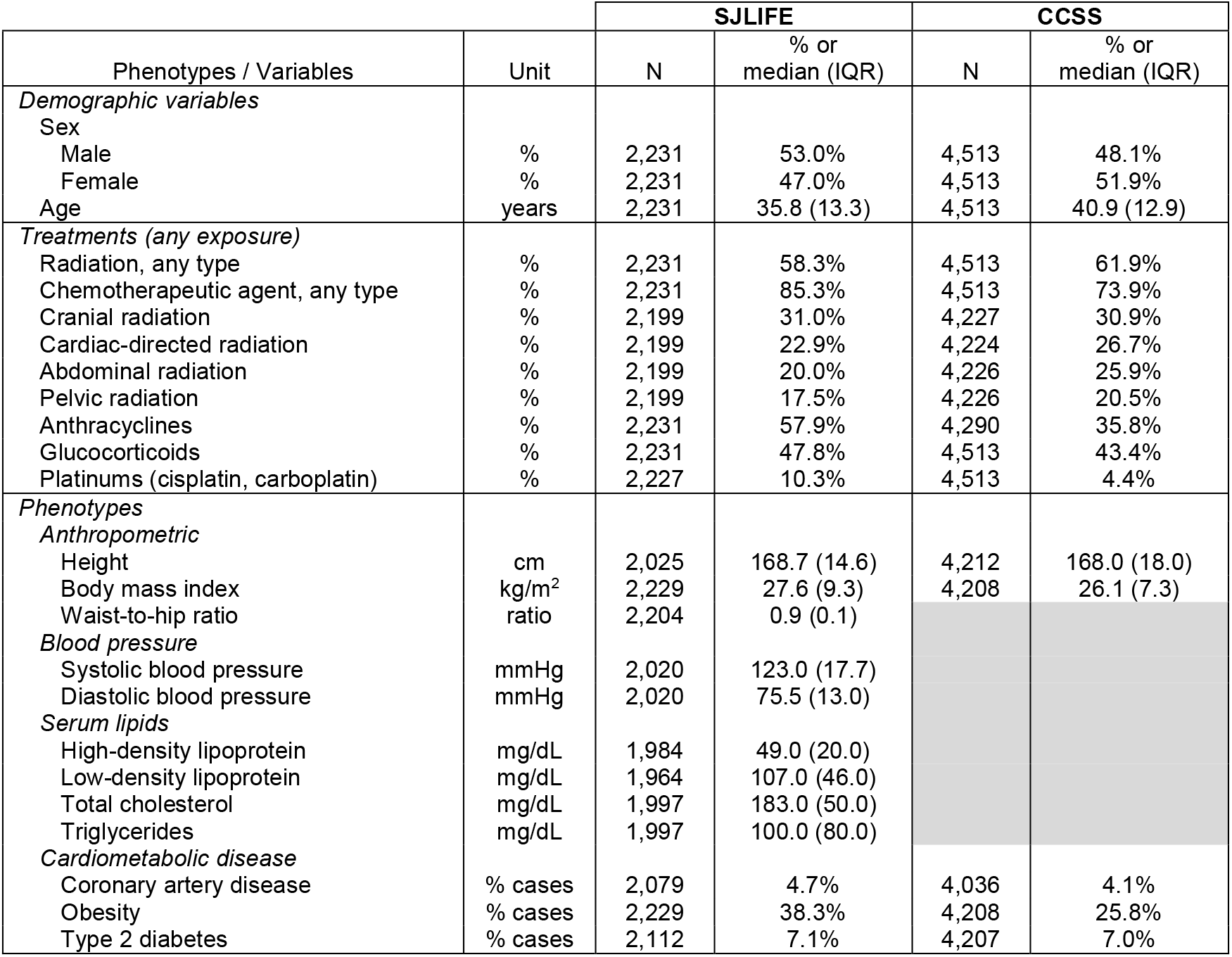
Descriptive statistics for phenotypes, treatments, and demographic variables in SJLIFE

|  |  | SJLIFE |  | CCSS |  |
| --- | --- | --- | --- | --- | --- |
| Phenotypes / Variables | Unit | N | % or<br>median (IQR) | N | % or<br>median (IQR) |
| *Demographic variables* |  |  |  |  |  |
| Sex |  |  |  |  |  |
| Male | % | 2,231 | 53.0% | 4,513 | 48.1% |
| Female | % | 2,231 | 47.0% | 4,513 | 51.9% |
| Age | years | 2,231 | 35.8 (13.3) | 4,513 | 40.9 (12.9) |
| *Treatments (any exposure)* |  |  |  |  |  |
| Radiation, any type | % | 2,231 | 58.3% | 4,513 | 61.9% |
| Chemotherapeutic agent, any type | % | 2,231 | 85.3% | 4,513 | 73.9% |
| Cranial radiation | % | 2,199 | 31.0% | 4,227 | 30.9% |
| Cardiac-directed radiation | % | 2,199 | 22.9% | 4,224 | 26.7% |
| Abdominal radiation | % | 2,199 | 20.0% | 4,226 | 25.9% |
| Pelvic radiation | % | 2,199 | 17.5% | 4,226 | 20.5% |
| Anthracyclines | % | 2,231 | 57.9% | 4,290 | 35.8% |
| Glucocorticoids | % | 2,231 | 47.8% | 4,513 | 43.4% |
| Platinums (cisplatin, carboplatin) | % | 2,227 | 10.3% | 4,513 | 4.4% |
| *Phenotypes* |  |  |  |  |  |
| *Anthropometric* |  |  |  |  |  |
| Height | cm | 2,025 | 168.7 (14.6) | 4,212 | 168.0 (18.0) |
| Body mass index | kg/m<sup>2</sup> | 2,229 | 27.6 (9.3) | 4,208 | 26.1 (7.3) |
| Waist-to-hip ratio | ratio | 2,204 | 0.9 (0.1) |  |  |
| *Blood pressure* |  |  |  |  |  |
| Systolic blood pressure | mmHg | 2,020 | 123.0 (17.7) |  |  |
| Diastolic blood pressure | mmHg | 2,020 | 75.5 (13.0) |  |  |
| *Serum lipids* |  |  |  |  |  |
| High-density lipoprotein | mg/dL | 1,984 | 49.0 (20.0) |  |  |
| Low-density lipoprotein | mg/dL | 1,964 | 107.0 (46.0) |  |  |
| Total cholesterol | mg/dL | 1,997 | 183.0 (50.0) |  |  |
| Triglycerides | mg/dL | 1,997 | 100.0 (80.0) |  |  |
| *Cardiometabolic disease* |  |  |  |  |  |
| Coronary artery disease | % cases | 2,079 | 4.7% | 4,036 | 4.1% |
| Obesity | % cases | 2,229 | 38.3% | 4,208 | 25.8% |
| Type 2 diabetes | % cases | 2,112 | 7.1% | 4,207 | 7.0% |

All meta-GWAS hits that were replicated in SJLIFE (*P*<0.05, with same directions of effect in literature) are listed in Supplementary Table 1. The results of the meta-GWAS hit replication enrichment analysis in SJLIFE are summarized in Figure 2 and Supplementary Table 2. Of the 1,376 meta-GWAS hits, we expected to replicate ~279 SNP-phenotype associations across all phenotypes, based on power calculations for replication with SJLIFE sample sizes and SNP EAFs. We replicated only 189 SNP-phenotype associations (replication rate=13.7%; 189/1,376 tested) with models adhering to reference GWAS, and 185 SNP-phenotype associations (replication rate=13.4%; 185/1,376 tested) after adjusting for additional covariates relevant to childhood cancer survivors (i.e., cancer treatment exposures, Table 1). The Replication Enrichment Ratio (RER), or the ratio of observed-to-expected meta-GWAS hit replication frequencies, across all 12 phenotypes was 0.68 (95% CI: 0.60-0.77, *P*=2.4×10^−9^) using models adjusting for reference GWAS covariates only, suggesting that the overall number of meta-GWAS hit replications observed in SJLIFE was significantly less than expected. Significant replication depletion was also observed across all phenotypes using models adjusting for additional covariates relevant to survivors (RER=0.66, 95% CI: 0.58-0.76, *P*=4.1×10^−10^). While three phenotypes (WHR, T2D, TG) showed no evidence of replication depletion (RER>1), the remaining nine phenotypes had either significant depletions of meta-GWAS hit replications (RER<1 and *P*<0.05 for height, BMI, DBP, and obesity) or suggestive evidence of replication depletions (RER<1 and *P*<0.2 for SBP, HDL, LDL, TC, CAD).

**Figure 2:**
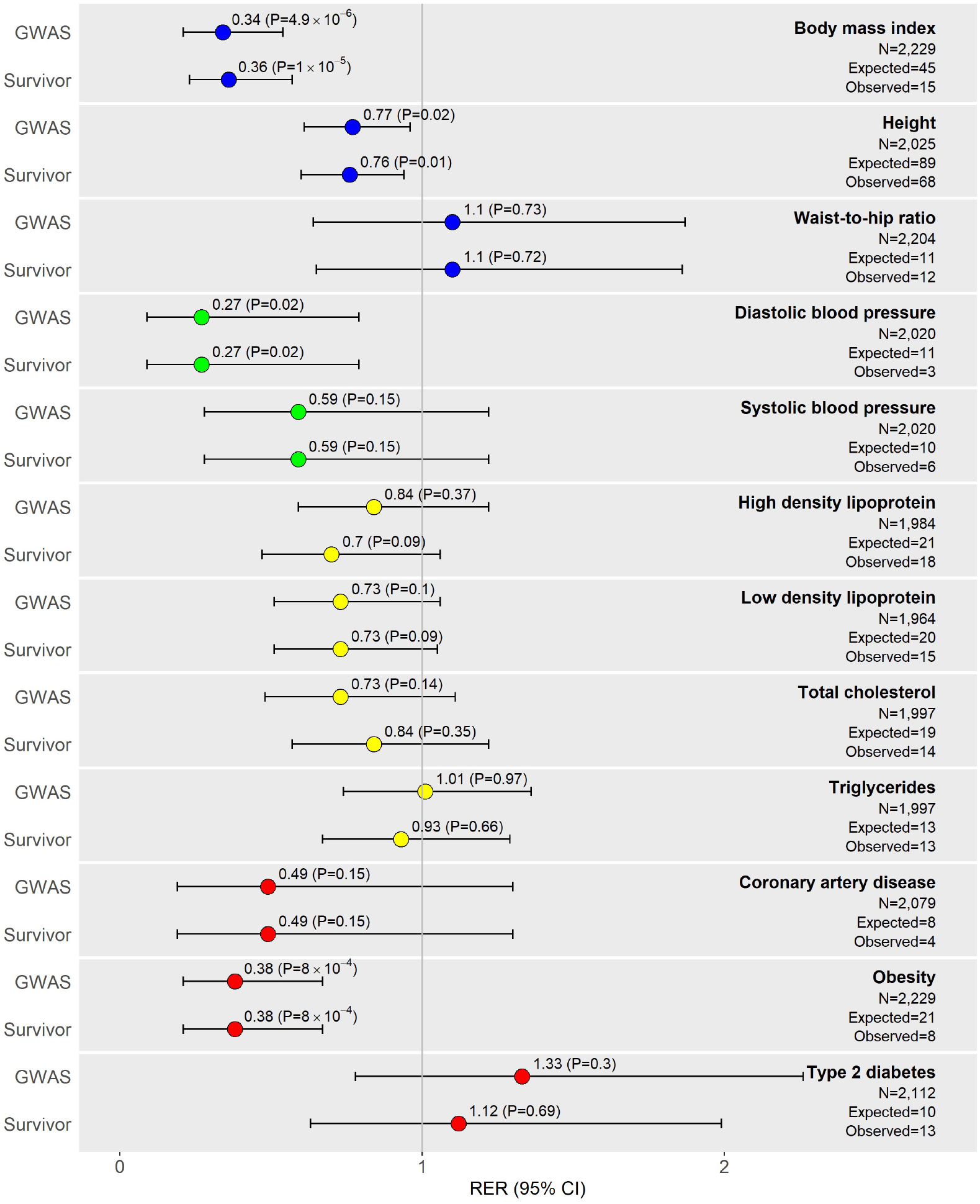
Plots of replication enrichment ratios (RERs) and respective 95% confidence intervals by phenotype in SJLIFE. RERs left of the vertical line corresponding to a RER equal to 1 suggest meta-GWAS hit replication depletion, i.e., observations of fewer replications of meta-GWAS hits than expected. RERs considering adjustment covariates under two different models are presented for each phenotype: (1) covariates adhering to reference GWAS (“GWAS”), and (2) GWAS covariates along with covariates considered in childhood cancer survivor populations (“Survivor”). Phenotype RERs are color-coded by similarity: anthropometric (blue); blood pressure (green); lipid (yellow), and cardiometabolic disease (red). The observed numbers of replications included in the figure are under the “GWAS” model. The expected numbers of replications are estimated by the sum of the power to replicate each SNP-phenotype association assuming observed SNP effect allele frequencies, the cohort sample size, an additive genetic inheritance model, α=0.05, and effect sizes in reference meta-GWAS.

We explored alternative definitions of meta-GWAS hit replication in SJLIFE. First, we examined an “extended” replication strategy, under the possible but unlikely scenario that all SNPs involved in the 1,187 non-replicated robust meta-GWAS hits are weak representatives for nearby causal variants, but are in high linkage disequilibrium (LD) with causal variants in the same LD block. We re-tested non-replicated meta-GWAS hits using best SNP proxies for reported index SNPs, where best proxies were defined as SNPs in high LD with the index SNP (r^2^>0.8 in the 1000 Genomes^15^ European reference population or 1000G EUR) likely to fall in the same LD block (i.e., within a 5-kb window, based on median LD block sizes of ~2.5 kb reported in 1000G EUR^16^). While we re-tested 812 non-replicated SNP associations with at least one plausible proxy (median=3 proxies per index SNP), this added only 12 additional meta-GWAS hit replications (overall RER=0.72, 95% CI: 0.64-0.82, *P*=2.2×10^−7^) (Supplementary Table 3). We also assessed replication rates for a set of independent SNP-phenotype associations by limiting the SNP set to those with the highest EAF in SJLIFE among clusters of SNPs in high LD (r^2^>0.8, 500-kb window in 1000G EUR) for each phenotype, in order to avoid bias in replication rate estimates due to clusters of SNPs in high LD. The same nine phenotypes as our primary analysis continued to show significant or suggestive replication depletion using the pruned SNP-phenotype associations (Supplementary Table 4). Finally, we examined replications of meta-GWAS hits under strict replication *P*-value thresholds corrected for multiple testing. While replication of ~55 SNP-phenotype associations were expected under Bonferroni-corrected *P*-value thresholds, only 25 SNP-phenotype associations were replicated, most of which were related to BMI/obesity or blood lipid phenotypes (Supplementary Table 5).

### Replicating meta-GWAS hits in childhood cancer survivors in CCSS

To assess our findings from SJLIFE in an independent cohort, we conducted a second analysis in survivors from the Childhood Cancer Survivor Study (CCSS). We examined five self-reported phenotypes available in CCSS that corresponded with our SJLIFE analysis (height, BMI, CAD, obesity, and T2D) in 4,513 survivors with high-quality imputed genotype data (loci with imputation quality score r^2^>0.8, see Methods). Descriptive statistics for the CCSS study sample are provided in Table 2. Similar to SJLIFE, most CCSS survivors had been exposed to at least one type of chemotherapeutic agent (73.9%) or radiotherapy (61.9%). Under power calculations for replication with CCSS sample sizes and EAFs, we expected to replicate ~253 meta-GWAS hits. A total of 135 SNP-phenotype associations were successfully replicated in CCSS survivors with complete genotype, phenotype, and covariate data (up to N=4,212) using models consistent with reference GWAS. All five phenotypes showed significant (*P*<0.05) or suggestive (*P*<.2) meta-GWAS hit replication depletions than expected (Figure 3, Supplementary Table 2), contributing to an overall RER of 0.53 (P=1.1×10^−16^) using models adhering to reference GWAS.

**Figure 3:**
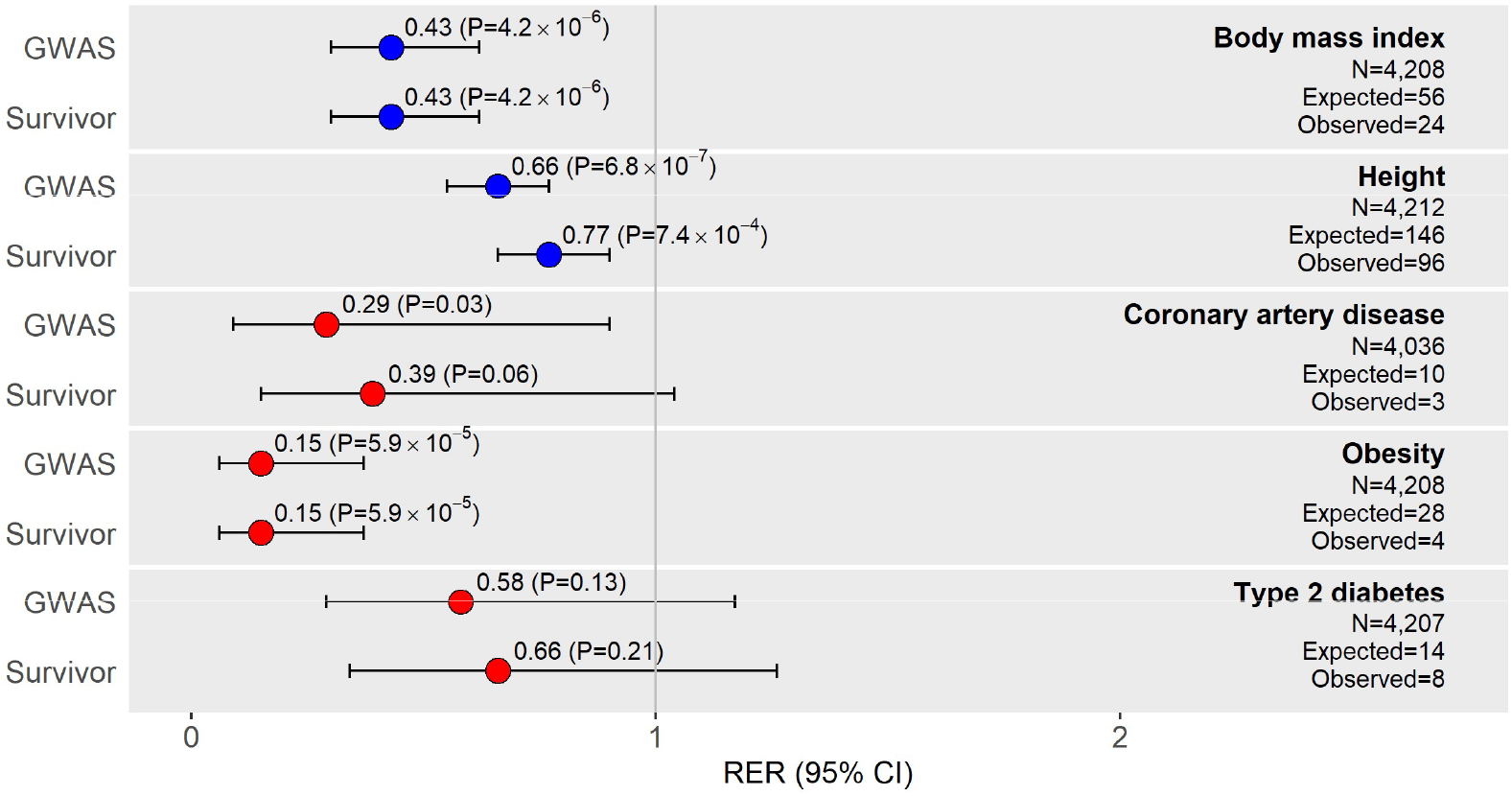
Plots of replication enrichment ratios (RERs) and respective 95% confidence intervals by phenotype in CCSS. RERs left of the vertical line corresponding to a RER equal to 1 suggest meta-GWAS hit replication depletion, i.e., observations of fewer replications of meta-GWAS hits than expected. Phenotype RERs are color-coded by similarity: anthropometric (blue) and cardiometabolic disease (red). The observed numbers of replications included in the figure are under the “GWAS” model.

### Treatments for pediatric cancer and meta-GWAS hit replication depletions in SJLIFE survivors

We considered whether factors specific to childhood cancer survivors, i.e., exposure to cancer treatments, could “disrupt” robust genetic associations reported in the general population. For the nine phenotypes that showed evidence of meta-GWAS hit replication depletion in SJLIFE (RER<1), we estimated RERs in survivor subgroups stratified by treatment exposure, where treatment exposure was defined as any exposure to therapeutic agents for pediatric cancer associated with the phenotype of interest (Table 1). We hypothesized that if cancer treatments contribute to phenotypic variation and obscure replications of meta-GWAS hits in survivors, we would not only observe replication depletion in treatment-exposed subgroups, but greater replication depletion in treatment-exposed subgroups than in treatment-unexposed subgroups.

We found evidence of replication depletion in treatment-exposed survivor subgroups for seven phenotypes: the height, BMI, TC, obesity, and DBP phenotypes showed significant (*P*<0.05) replication depletion, while CAD and LDL phenotypes showed suggestive (*P*<0.2) replication depletion. Among these seven phenotypes, CAD, height, LDL, TC, and DBP showed stronger evidence of replication depletion in treatment-exposed subgroups compared to treatment-unexposed subgroups (i.e., smaller RERs in treatment-exposed subgroups; Figure 4). Specifically, CAD, height, LDL, and TC also had the greatest changes in adjusted R^2^ (>1%) and the strongest treatment likelihood ratio test *P*-values (*P*<1×10^−7^) when comparing clinical models with and without the relevant treatments, suggesting that replication depletions in meta-GWAS hits are exacerbated in survivors when treatments have greater contributions to the phenotype risk.

**Figure 4:**
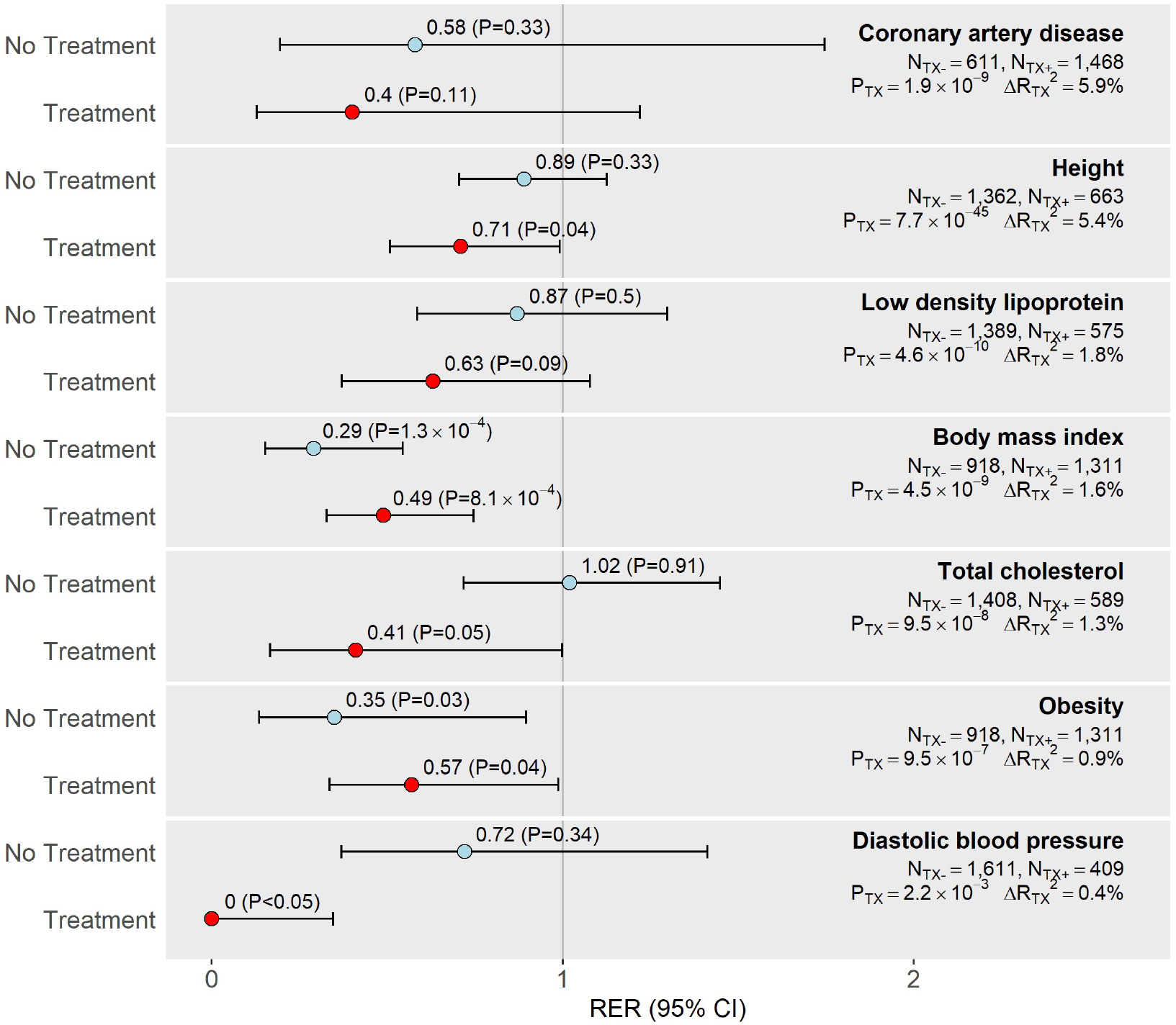
Plots of phenotype-specific replication enrichment ratios (RERs) and respective 95% confidence intervals in samples unexposed to treatments (“No Treatment”) and exposed to treatments (“Treatment”). Treatments were defined as cancer treatments associated with phenotypes. Phenotypes with any evidence of replication depletion (RER<1) in our main analysis that showed either significant (*P*<0.05) or suggestive (*P*<0.2) replication depletion in treatment-exposed samples are included in this figure. Sample sizes by exposure strata (N_TX−_, No Treatment; N_TX+_, Treatment) are provided, as well as likelihood ratio test *P*-values representing treatment associations with phenotypes (P_TX_) and changes in adjusted R^2^ (ΔRTX^2^) after removing treatment variables from clinical models. Phenotypes are ordered by ΔRTX^2^ values, with larger ΔRTX^2^ values reflecting greater treatment influence on phenotype variation.

### Differences in functional/epigenetic annotations for replicated and non-replicated meta-GWAS hits

We speculated that meta-GWAS SNPs with replicated phenotype associations in survivors could have functional/epigenetic annotation enrichments that may distinguish them from SNPs with non-replicated associations. Using publicly available bioinformatics data from GTEx^17^ and the Roadmap Epigenomics Consortium^18^ for functional/epigenetic annotation, we compared the set of 170 SNPs with at least one replicated association with the 12 phenotypes (“replicated SNPs”) against the set of 1,061 SNPs without any replicated associations (“non-replicated SNPs”) from our main analysis in SJLIFE. Similar proportions of replicated and non-replicated SNPs were mapped to RefSeq^19^ gene bodies (57.1% vs. 58.7%, respectively; *P*=0.74). Using GTEx^17^ to examine expression quantitative trait loci (*cis*-eQTL) enrichment, replicated SNPs had greater odds of being a *cis*-eQTL SNP (FDR≤0.05) in adipose and liver tissues than non-replicated SNPs (nominal *P*<0.05, Supplementary Table 6). Top 15-state ChromHMM^18^ enhancer and promoter chromatin state annotation enrichments revealed that replicated SNPs also had greater odds of overlapping enhancer chromatin states in cell/tissue types related to the kidney, adipose, gut and obesity-linked brain structures (nominal *P*<0.05, Supplementary Table 7). We also assessed top Reactome^20^ biological pathway enrichments for non-overlapping genes mapped to replicated and non-replicated SNPs against all other genes in human genome (Supplementary Figure 5). For the 79 genes that corresponded with the replicated SNPs, the lead biological pathway enrichments (FDR<0.10) were specific to cardiometabolic phenotypes, i.e., plasma lipoprotein metabolism is connected to serum lipid traits; elastic fiber assembly is related to arterial wall formation and cardiovascular phenotypes; PPARalpha-mediated lipid metabolism is linked to metabolic phenotypes. To contrast, the vast majority of lead biological pathway enrichments (FDR <0.10) for the 466 genes mapped to non-replicated SNPs were related to signal transduction.

### Treatment-DNA methylation patterns and non-replicated meta-GWAS hits in SJLIFE

We used BIOS Consortium (BIOS QTL^21^) methylation quantitative trait loci (meQTLs) as a reference resource for ancillary DNA methylation analyses. BIOS QTL includes samples from the Lifelines Cohort Study, which recently reported high meta-GWAS hit replication rates (median=84%) across 32 phenotypes^2^. Whole blood *cis*-meQTLs (≤250 kb between SNP and CpG) from BIOS QTL for any of the 1,231 meta-GWAS SNPs of interest (FDR<0.05) were regarded as established phenotype-variant-associated *cis*-meQTLs in the general population. Most meta-GWAS SNPs examined in our main analysis (87.5%, 1,077 SNPs) were mapped to at least one established *cis*-meQTL (Supplementary Table 8).

First, we assessed whether established *cis*-meQTLs in the general population (BIOS QTL) could be generalized to childhood cancer survivors using experimental blood-derived methylome and genotype data from 236 SJLIFE survivors. Despite the small sample size, we successfully validated 5,651 established *cis*-meQTLs for the meta-GWAS SNPs of interest (40.6%; 13,930 tested) in SJLIFE, where validation was defined by SNP-CpG methylation associations with *P*<0.05 and the same directions of association as reported in BIOS QTL. We further evaluated whether SJLIFE-validated *cis*-meQTLs could be differentiated by their relationships to SNPs with successful or failed replications in survivors. We discovered that non-replicated SNPs had greater odds of being *cis*-meQTLs than replicated SNPs (OR=1.66, *P*=0.02, Supplementary Table 9).

Next, we investigated the involvement of *cis*-meQTLs in meta-GWAS hit replications in SJLIFE by considering whether replications were affected by childhood cancer treatments. Specifically, we compared 48 “treatment-sensitive” meta-GWAS SNPs that showed replicated assocations only in the treatment-unexposed subgroup, i.e., in survivors that are more similar to the general population, and 66 “treatment-insensitive” meta-GWAS SNPs with robust replications, i.e., replicated in both treatment-unexposed and treatment-exposed subgroups. We found greater enrichment for SJLIFE-validated *cis*-meQTLs among treatment-sensitive SNPs (38/42, 90.5%) compared to treatment-insensitive SNPs (37/57, 64.9%; OR=5.06, *P*=4.1×10^−3^, Supplementary Table 9), suggesting that SNPs with phenotype association replications that were perturbed by treatment exposures in survivors were more likely to involve *cis*-meQTL mechanisms than SNPs with robust replications.

We then explored whether non-replicated meta-GWAS hits in survivors could be attributed to treatment-related disruptions of *cis*-meQTL profiles. We hypothesized that survivors’ exposures to treatments that counter the direction of CpG methylation by a meta-GWAS SNP would reduce the likelihood of replication for the corresponding SNP-phenotype association in survivors. We measured treatment-related disruptions of *cis*-meQTL profiles by counting the frequency of discordance in the direction of methylation at a CpG site in BIOS QTL for a meta-GWAS SNP and the direction of methylation at the same CpG site for exposure to a specific childhood cancer treatment. We split the 4,153 CpG sites linked to the 5,561 SJLIFE-validated *cis*-meQTLs between replicated and non-replicated SNPs, i.e., 549 “replicated CpGs” versus 3,604 “non-replicated CpGs”, respectively. We examined different radiation therapy (RT) and chemotherapeutic exposures (Supplementary Table 10). Non-replicated CpGs were enriched for directionally discordant SNP-methylation and treatment-methylation associations for multiple treatment types relative to the replicated CpGs (Supplementary Table 11). The non-replicated CpGs showed the strongest enrichment for directionally discordant methylation associations for pelvic RT, with ~54% of non-replicated CpGs bearing directionally discordant methylation associations in contrast to ~29% of replicated CpGs (OR=2.90, P=8.7×10^−4^). The non-replicated CpGs were also significantly enriched for directionally discordant associations for chest RT (OR=2.70, P=5.3×10^−4^) and modestly enriched for abdominal RT (OR=1.91, P=0.06).

We illustrate these results by describing the failed replication of the T2D risk variant rs1552224 (chr11:72722053, GRCh38) in SJLIFE survivors as an example. Multiple meta-GWAS have linked the A allele of rs1552224 with increased T2D risk^22,23^. However, this association was not replicated among survivors exposed to abdominal or pelvic RT, but was replicated in survivors without these RT exposures (Supplementary Table 12). Figure 5 demonstrates how abdominal/pelvic RT can obscure the replication of the rs1552224 – T2D risk association in survivors by disrupting *cis*-meQTL effects on T2D risk in the general population. The strongest *cis*-meQTL effect for rs1552224 was reported at cg04827223 in BIOS QTL (assessed allele=A, Z=34.8, *P*=6.0×10^−266^) and was validated in SJLIFE (β=0.12, *P*=3.7×10^−4^). Figure 5a shows increasing A allele dose for rs1552224 corresponds with increases in methylation at cg04827223 and T2D risk in survivors without exposures to abdominal/pelvic RT, consistent with the general population. But in survivors with increasing doses of abdominal/pelvic RT, increasing A allele dose for rs1552224 does not change methylation at cg04827223 or T2D risk (Figure 5b, 5c), which reflects the inverse relationships between methylation levels at cg04827223 and pelvic (β=−4.0×10^−6^, *P*=0.03) and abdominal RT (β =−3.4×10^−6^, *P*=0.06) dose observed in SJLIFE.

**Figure 5:**
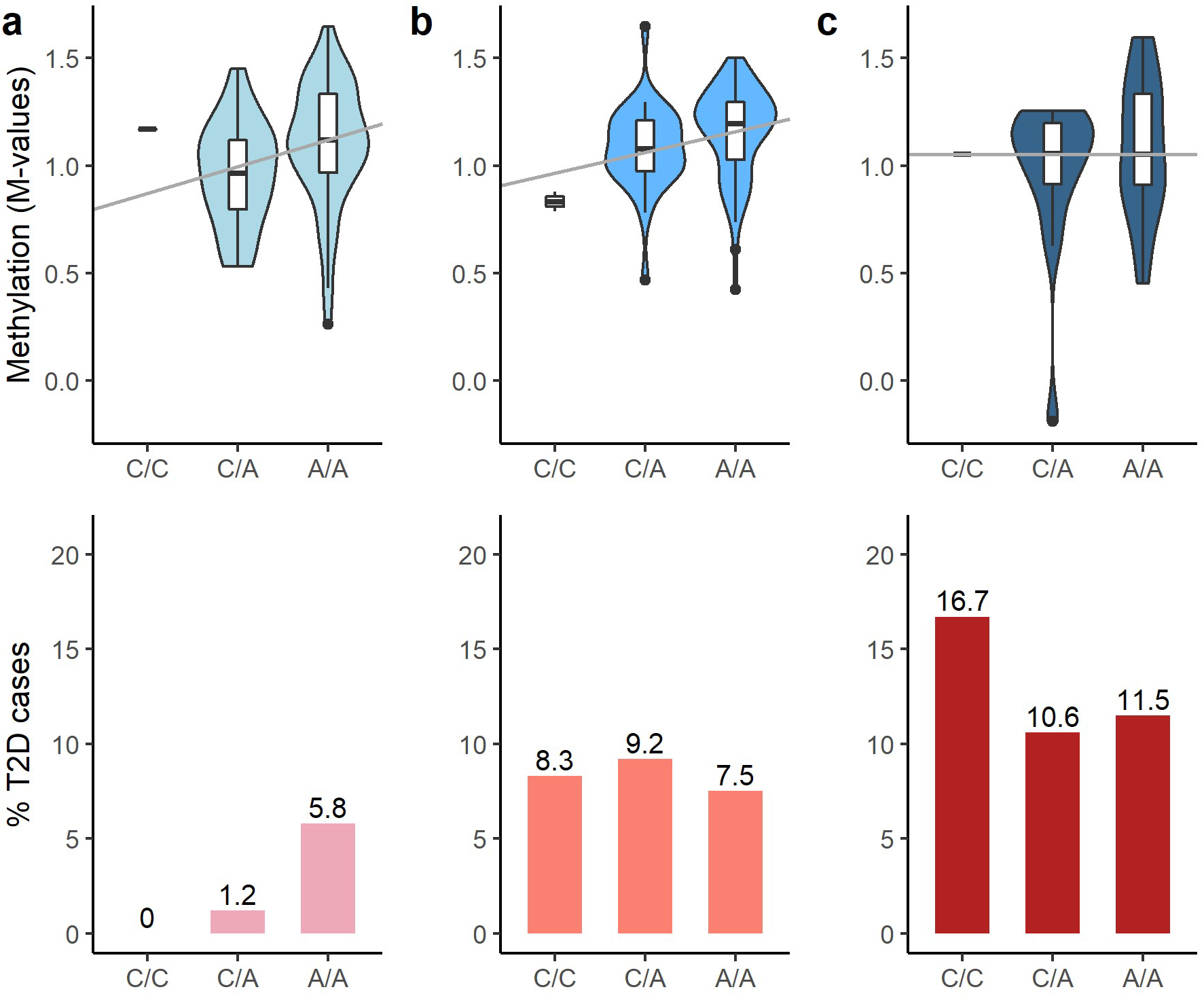
DNA methylation levels at cg04827223 and percentage of T2D cases by genotype classes for rs1552224 in SJLIFE survivor subgroups with no (a), low-to-moderate (b), and high doses (c) of abdominal or pelvic radiation therapy (RT). No RT dose was defined as 0 Gy, low-to-moderate RT dose was defined by >0 to <20 Gy, and high dose was defined by ≥20 Gy. The upper panels show the observed methylation level relationships with the SNP at the cg04827223 CpG site in the SJLIFE subset with methylome and genotype data (N=236); boxes represent the median and interquartile range (IQR), with whiskers extending from the first or third quartile to 1.5 times the IQR. Methylation level trend by allele dose is shown with median regression lines. Genotype frequencies in this SJLIFE subset were as follows: 1.8% (C/C), 30.8% (C/A), and 67.4% (A/A). The lower panels show the percentage (%) of T2D cases by genotype in SJLIFE survivors in the main analysis (N=2,112), with the following genotype frequencies: 1.9% (C/C), 26.9% (C/A), and 71.2% (A/A).

## DISCUSSION

There is growing interest in leveraging knowledge of established meta-GWAS hits though polygenic risk scores (PRS) in specialized clinical populations such as childhood cancer survivors^24^. The suitability of translating this knowledge to such populations, however, depends on the generalizability of general population SNP associations to the clinical population of interest. We evaluated the generalizability of 1,376 SNP associations reported in 46 selected meta-GWAS for 12 anthropometric and cardiometabolic phenotypes in a large cohort of adult survivors of pediatric cancer in SJLIFE using genotypes from whole genome sequencing and clinically ascertained phenotypes. Significantly fewer than expected robust meta-GWAS hits were replicated in SJLIFE survivors, with an observed-to-expected RER of 0.68 (*P*=2.4×10^−9^) across all phenotypes. Replication depletion was also observed in a secondary analysis of five comparable phenotypes in an independent cohort of survivors from CCSS. These results suggest that advances in genetic risk prediction (and opportunities for targeted intervention) in vulnerable clinical populations like childhood cancer survivors may ultimately lag behind the general population, and highlight the need for novel genetic association studies in diverse populations.

Given that the meta-GWAS hits we tested were robust findings in the general population, i.e., were genome-wide significant (*P*<5×10^−8^) and compiled from large meta-GWAS (>10,000 participants), and accompanied by replication, complementary functional annotation, and even experimental validation studies, the limited generalizability of these genetic associations to survivors is unexpected. For comparison, one of the largest recent studies of the generalizability of European-derived GWAS hits in a non-European, multi-ancestral population (N=49,839) observed a more reasonable ~42% replication rate (*P*<0.05 threshold) across 22 complex continuous phenotypes^25^, despite the accumulating evidence for the poorer predictive accuracy of European-derived PRS in non-Europeans^1^. Discovering that these meta-GWAS hits may only be partially generalizable to survivors is unlikely to be attributable to the methods we employed: we tested associations between measured (not imputed) index SNPs and clinically ascertained phenotypes; we restricted our analyses to survivors of European ancestry; we observed high correspondence between EAFs in SJLIFE and the reference literature; and replication depletion was evaluated accounting for the expected probability of replication based on our sample size.

We further investigated the possible but unlikely scenario that non-replications could be primarily due to testing index SNPs that were poor representatives for SNPs causal for phenotype in the same LD block, or non-replication bias due to highly correlated clusters of non-replicating SNPs. These ancillary analyses, along with our analysis of five corresponding phenotypes in a second cohort of survivors in CCSS, corroborate that some of these meta-GWAS hits do not apply to survivors. This analysis is among the first to provide evidence towards a hypothesis described in a recent review of the transferability of PRS across populations, specifically that the generalizability of PRS may also be limited in cohorts with differential environmental exposures^1^.

Recent studies have demonstrated that ionizing radiation can induce persistent changes in DNA methylation in cells/tissues targeted by radiation that are dose-dependent^26–30^. Chemotherapies, e.g., cisplatin^31^ and carboplatin^32^, have also been linked to differential methylation of genes involved in cell cycle regulation and DNA repair. In this study, we discovered when cancer treatments had greater contributions to phenotype risk, greater replication depletions than expected were observed in treatment-exposed survivor subgroups. Therefore, we assessed whether treatment-related DNA methylation could potentially “disrupt” robust SNP-phenotype relationships reported in the general population among survivors. We found that non-replicated SNPs were significantly enriched for SNPs with *cis*-meQTLs reported in BIOS QTL that were also validated in a subset of SJLIFE survivors. Furthermore, we discovered a ~5-fold enrichment (*P*=4.1×10^−3^) of validated *cis*-meQTL SNPs among SNPs with replications perturbed by treatments in survivors compared to SNPs that were robustly replicated in survivors. Lastly, enrichments of “disruptive” or directionally discordant methylation associations for chest (OR=2.70, P=5.3×10^−4^), pelvic (OR=2.90, P=8.7×10^−4^), and abdominal (OR=1.91, P=0.06) RT among CpGs linked to meta-GWAS SNPs that failed to replicate in SJLIFE survivors were observed. Notably, chronic hematological toxicity has been well-documented for RT to the chest, pelvic, and abdominal fields due to the volume of active bone marrow in these regions^33^, which suggests the DNA methylation patterns we see in the blood-derived methylome data are plausibly related to these RT exposures. Taken together, these results suggest cancer treatments (particularly RT), may disrupt DNA methylation patterns at genomic sites linked to some disease-/trait-associated variants and interfere with their generalizability to survivors.

The main limitation of this analysis was the relatively small sample sizes of the survivor cohorts. Our analysis had limited power to detect some SNP-phenotype replications (especially those with small effect sizes), but we estimated the expected number of replications given available power accounting for sample size, reported effect sizes, and sample EAFs and used these estimates to compare observed and expected replication rates. We also performed a secondary analysis of meta-GWAS hit replications in the CCSS cohort which was nearly double the size of the SJLIFE cohort and saw stronger evidence of replication depletions. Another limitation was that we could not combine CCSS and SJLIFE cohorts for all 12 phenotypes, since all phenotypes in CCSS are self-reported. Lastly, interpretations of our analyses of SNP and treatment associations with cross-sectional whole blood DNA methylation measurements have several limitations. We were only able to evaluate DNA methylation associations in a small sample of survivors (N=236); however, we did observe a high (~41%) validation rate for established *cis*-meQTLs (FDR<0.05) reported by BIOS QTL. Similar to the limitations reported in other analyses of DNA methylation associations, we cannot ascertain the extent to which methylation levels at the selected CpGs truly contribute to phenotype variation, or that methylation associations with treatments are strictly attributable to our factor of interest (treatments) versus some other related factor with potential effects on DNA methylation (e.g., primary cancer diagnosis). In addition, evaluating associations between treatments and gene expression levels linked to these CpG sites would be a necessary first step to determine how treatment-related changes in DNA methylation disrupt SNP-phenotype associations. Despite these limitations, our preliminary analyses of DNA methylation in survivors have specific strengths: cumulative prior exposures to RT and chemotherapy are well-documented in our sample, and our analyses only examine established meta-GWAS variants and *cis-*meQTLs.

In summary, we have shown that robust meta-GWAS SNP hits that were observed in general populations for a range of cardiometabolic phenotypes are only partially generalizable to childhood cancer survivor cohorts. Methodologies and applications that rely on established meta-GWAS hits from the general population to predict or clinically surveil some cardiometabolic outcomes or traits may have limited utility in survivors. A plausible explanation for the partial generalizability of robust meta-GWAS hits in survivors is that cancer treatment exposures obscure some genetic associations through epigenetic alterations such as DNA methylation. This phenomenon may also apply to other clinical populations.

## METHODS

### Compiling SNP associations with complex traits and diseases

We selected 12 complex traits and diseases that were: (a) related to cardiovascular and metabolic disease; (b) measured or clinically ascertained during SJLIFE study visits; and (c) examined in at least one recent (i.e., published after 01/01/2008) meta-GWAS with >10,000 participants of European ancestry. To identify genetic associations for our replication analysis, we searched all reports available in the NHGRI-EBI GWAS Catalog^14^ published between 1/1/2008 – 11/20/2017 and retained any meta-analysis based on the following reference literature selection criteria: (1) study is relevant to the phenotype and the association testing method of interest (i.e., no SNP interaction or gene-environment interaction association testing); (2) study was performed in predominantly European cohort(s); (3) study included a replication analysis; and (4) study had discovery and/or replication sample size(s) with at least 10,000 participants (Figure 1). We reviewed the compiled literature to confirm the set of “index SNPs” for replication testing, i.e., published SNPs with genome-wide significant associations (*P*<5×10^−8^), and their respective effect sizes, *P*-values, and effect alleles. Effect allele frequencies (EAFs) and standard errors were recorded when available. Reported effect sizes and *P*-values for a published SNP association were taken from the combined analysis of discovery and replication samples; if a combined analysis was not available, effect sizes were taken from the replication analysis and *P*-values were taken from the discovery analysis. When necessary, we transformed effect sizes reported in different units across papers for comparability.

### Description of study cohorts

This study was approved by the Institutional Review Boards at St. Jude Children’s Research Hospital (SJCRH; Memphis, TN) and all participating study centers. All participants in this study provided informed consent. Brief descriptions of the two cohorts included in our study are provided below. Additional details regarding phenotype-specific analyses applied in both cohorts, including reference GWAS-informed definitions for phenotypes, adjustment covariates, and participant exclusion criteria, along with survivor-specific factors, are provided in Table 1.

#### SJLIFE cohort

Initiated in 2007, the St. Jude Lifetime Cohort Study^34^ (SJLIFE) is an ongoing retrospective cohort study dedicated to the longitudinal study of a wide-ranging set of health outcomes in survivors treated for pediatric cancer at SJCRH. The details of this study have been described previously^34^. In brief, eligibility criteria include treatment for pediatric cancer at SJCRH and ≥5 years survival since diagnosis. Participants included in the current study were ≥18 years of age, had no history of allogeneic stem cell transplantation, participated in specimen biobanking, and completed at least one SJCRH study visit as of the June 30, 2015 freeze date.

SJCRH study visits include medical evaluations (with core laboratory/diagnostic studies), assessments of self-reported outcomes, and examinations of neurocognitive function and physical performance. Data for demographics, treatments (chemotherapeutic agent cumulative dosages; field/doses of radiation therapy; surgical interventions), and primary cancer diagnosis were obtained from medical record review. Medication use was self-reported as a part of the health and behavior questionnaires. All quantitative trait measurements used in this analysis were taken from the participant’s most recent SJLIFE study visit as of 06/30/2017. Height and weight were measured using a stadiometer and an electronic scale (Scale-Tronix, White Plains, NY); WHR circumferences were taken with a Gulick tape measure. BMI values were adjusted for amputation. Average systolic and diastolic blood pressure (SBP and DBP, respectively; mmHg) values for participants with at least two measurements taken with a calibrated sphygmomanometer after an initial 5-minute rest were used. Fasting blood lipids (mg/dL), including high-density lipoprotein (HDL), calculated low-density lipoprotein (LDL), total cholesterol (TC), and triglycerides (TG) were measured using an enzymatic spectrophotometric assay (Roche Diagnostics, Indianapolis, IN).

Coronary artery disease (CAD) and diabetes mellitus were clinically assessed and graded according to the SJCRH-modified NCI Common Terminology Criteria for Adverse Events (CTCAE) v4.03 classification system^35^. The CTCAE grades used to define cases were based on presence of symptoms and/or relevant medication use. For CAD, use of medications to treat angina symptoms or evidence of abnormal cardiac enzymes, angina and ischemic heart disease, myocardial infarction, percutaneous transluminal coronary angioplasty (PTCA), or coronary artery bypass grafting (CABG) was used to define cases. Participants with symptomatic diabetes or use of oral medications or insulin to treat diabetes were considered as diabetes mellitus cases; for this analysis, we treated all cases of diabetes mellitus as type 2 diabetes cases (T2D) given recent reports suggesting that at least 79% of cases in survivors can be classified as T2D^36^. Brief episodes of diabetes mellitus occurring immediately after treatment or pregnancy were excluded. Obesity was defined as BMI ≥30kg/m^2^, which was consistent with CTCAE-based obesity grades.

#### CCSS cohort

The Childhood Cancer Survivor Study^37^ (CCSS) is a retrospective cohort study of 5-year childhood cancer survivors with prospective follow-up. Descriptions for CCSS participant eligibility and study design have been published in detail elsewhere^38,39^. CCSS participants included in this analysis were <21 years of age at primary cancer diagnosis between January 1, 1970 and December 31, 1986, received treatment for pediatric cancer at one of 26 participating study institutions in North America, responded to at least one CCSS questionnaire covering demographics, health conditions, health-related behaviors and health care use; and provided a whole blood, saliva, or buccal sample for DNA sequencing.

All phenotypes assessed in CCSS (height, BMI, obesity, CAD, T2D) were self-reported or reported by family proxies for survivors who could not complete surveys, were deceased or <18 years old. For CAD and T2D phenotypes, questionnaire responses related to these conditions (including relevant medication use) were graded using CTCAE v4.03. Information related to chemotherapy, radiotherapy, and surgery was abstracted from medical records. Participants with height values above/below ±4 SD of the sample mean or improbable BMI values (<10, >100 kg/m^2^) were excluded from analyses. Exclusion criteria or covariates considered in analyses performed in SJLIFE that were not included in CCSS due to missing data included genetic conditions affecting height and hypothalamic-pituitary axis tumor history. Any exposure to glucocorticoids was used as a substitute for glucocorticoid cumulative dosages. All other exclusion criteria, adjustment covariates, and case/phenotype definitions were identical to those applied to the SJLIFE analysis.

### Genotype data

Our analysis was restricted to the common SNPs (≥1% EAF) reported to have a genome-wide significant association (*P*<5×10^−8^) with any of the selected phenotypes in the meta-GWAS that met our reference literature selection criteria (i.e., index SNPs). We also considered best common SNP proxies, defined as SNPs in high LD with corresponding index SNPs in the European 1000 Genomes^15^ (1000G EUR) populations (minimum r^2^ =0.8) likely to fall in the same LD block. Descriptions for collecting and processing genotype data for each cohort are summarized below.

#### SJLIFE genotype data

The SJLIFE genotype data used in this analysis was collected as a part of larger effort to sequence whole genomes of SJLIFE participants^40^. Comprehensive details of DNA sample collection, extraction, sequencing, quality control, and variant mapping have been described previously^40,41^. Briefly, sequencing for 3,006 samples was completed at the HudsonAlpha Institute for Biotechnology Genomic Services Laboratory (Huntsville, AL) using the Illumina HiSeq X10 platform to yield 150 base pair paired-end reads with an average coverage per sample of 36.8X. Whole exome data from survivors (coverage >20x) sequenced by the SJCRH Department of Computational Biology was used to assess the validity of coding variants. Sequenced data was aligned to the GRCh38 human reference using BWA-ALN v0.7.12^42^. Variant calls were processed with GATK v3.4.0^43^ and BCFtools^44^. PLINK v1.90b^45^ and VCFtools v0.1.13^46^ were used to perform additional quality control, applying the following sample exclusion criteria: excess missingness (≥5%), cryptic relatedness (pi-hat>0.25), and excess heterozygosity (>3 SD). Variants with Hardy Weinberg Equilibrium (HWE) *P*<1×10^−10^ and >10% missingness across samples were removed, leaving approximately 84.3 million autosomal single nucleotide variants (SNVs) and small insertions and deletions (indels) in 2,986 samples. We then restricted our sample to the 2,364 participants that were identified as European (see *Ancestry* below).

#### CCSS genotype data

Details describing methods used to generate genotype data for the CCSS cohort can be found in previous papers^47,48^. To summarize, DNA was extracted from whole blood, saliva, or buccal samples and genotyped at the Cancer Genomics Research Laboratory of the National Cancer Institute (Bethesda, MD) using the Illumina HumanOmni5Exome array. Genotyping Module v1.9 (Illumina GenomeStudio software v2011.1) was used to call genotypes. The following per-sample exclusion criteria were applied: ≥8% missingness, heterozygosity of <0.11 or >0.16, X chromosome heterozygosity >5.0% for males or <20.0% for females, and identity-by-descent sharing >0.70. Genotypes were then imputed using Minimac3^49^ and the Haplotype Reference Consortium r1.1 reference panel for the 5,739 samples meeting quality control thresholds. After retaining 4,513 survivors of European ancestry (see *Ancestry* below) with no overlap with SJLIFE, downstream analyses excluded SNPs with minor allele frequency <1% and missingness >5% and only considered SNPs with high imputation quality (r^2^≥0.8).

#### Ancestry

Procedures to identify the ancestry of SJLIFE and CCSS samples have been described elsewhere^41,48^. Briefly, PLINK v1.90b was used to perform an EIGENSTRAT-based Principal Component Analysis^50^ for each cohort by combining the cohort samples with samples from 1000G global reference populations. Cohort samples with principal component scores within 3 SD of the means of the first two principal components in the 1000G EUR populations were of European ancestry.

### SJLIFE DNA methylation data

Whole blood DNA methylation was measured in 300 survivors in SJLIFE with a range of treatment histories with the Infinium MethylationEPIC Array (Illumina, San Diego, CA, USA) according to the manufacturer’s protocols. Genomic DNA (500 ng per sample; previously extracted for WGS) was treated with bisulfate using the Zymo EZ DNA Methylation Kit under the following thermos-cycling conditions: 16 cycles: 95°C for 30 sec, 50°C for 1 hour. Following bisulfite treatment, DNA samples were desulphonated, column purified, then eluted using 12 μl of elution buffer (Zymo Research). Bisulfite-converted DNA (4 μl) was then processed by following the Illumina Infinium Methylation Assay protocol which includes hybridization to MethylationEPIC BeadChips, single base extension assay, and staining and scanning using the Illumina HiScan system. The raw intensity data was exported from the Illumina Genome Studio Methylation Module as IDAT files for further downstream analysis.

Raw intensity data was processed with the “minfi” R package^51^, including sample and probe quality controls, background correction, and normalization. Probes were mapped to the GRCh38 build to identify and remove cross-reactive and non-specific probes. We eliminated samples with a low call rate (<95% probes with a detection *P* value <0.01) or sex discrepancies, along with probes located on sex chromosomes, with low detection rates (<95%), or with SNPs at CpG sites. A total of 689,742 high-quality probes were retained for 300 samples after preliminary quality control. Of the 15,481 probes in BIOS QTL contributing to significant *cis-*meQTLs with meta-GWAS SNPs of interest, 11,458 probes were available for the current study after quality control for the 236 participants of European ancestry with WGS data that were included in our main analysis.

### SNP-phenotype association testing and replication enrichment analysis

Statistical procedures to perform SNP-phenotype association testing and replication enrichment analysis were identical in SJLIFE and CCSS cohorts. Details are described below.

#### SNP-phenotype association testing

We conducted association tests for the reported genome-wide significant SNPs using phenotype definitions (i.e., units and transformations), exclusion criteria, and adjustment covariates that were consistent with the literature, along with factors relevant to childhood cancer survivors (Table 1). All regression coefficients, standard errors, and *P*-values were obtained with linear or logistic regression for quantitative traits or disease outcomes, respectively, using R v3.4.1. All association tests assumed an additive model of genetic inheritance. We used the first 10 principal components as covariates in all association analyses to account for population stratification. Measurements for adjustment covariates or data applied for phenotype transformations that were closest to the measurement or validation date of the trait/outcome were taken. SNP-phenotype associations with *P*-values <0.05 and the same direction of effect as the reference literature were considered as successful replications. While we also evaluated replications under trait-specific Bonferroni-corrected *P*-value thresholds, we regarded the *P*-value threshold of 5% as the primary definition for replication because all tested SNP associations were considered to be robust associations, i.e., published in large-scale meta-GWAS. In SJLIFE, we considered whether reported index SNPs were in high LD with potentially “causal” SNP candidates that would better capture the phenotype association at a given locus or LD block. To this end, we tested all best SNP proxies for non-replicated SNP associations, where best proxies for an index SNP were defined as SNPs in strong LD with the index SNP in the 1000G EUR populations (r^2^>0.8) within a 5-kb window of the index SNP (based on a median LD block size of ~2.5 kb^16^ in 1000G EUR). We also assessed observed versus expected replication rates for a pruned set of independent SNP-phenotype associations in SJLIFE given that non-replication rates from clusters of high-LD SNPs without replication signals could inflate replication depletions. Pruning entailed retaining the SNP with the highest EAF in SJLIFE among clusters of SNPs in high LD (r^2^>0.8, 500-kb window in 1000G EUR) for each phenotype.

#### Replication power and enrichment analysis

We used QUANTO v1.2.4^52^ to estimate the power for replicating each SNP association reported in the compiled literature with its respective phenotype in SJLIFE and CCSS. Power calculations assumed a 5% significance threshold (as well as a Bonferroni-corrected significance threshold in SJLIFE), phenotype-specific sample sizes, and an additive genetic model. Phenotype-specific power curves for our main analysis accounting for a range of effect allele frequencies and effect sizes are provided in Supplementary Figures 1–4. We used these power calculations to estimate the replication power for each SNP-phenotype association assuming the effect size reported in reference GWAS and the effect allele frequency observed in the survivor cohorts. We used the same procedure to also estimate replication power for each SNP-phenotype association in treatment-exposed and treatment-unexposed subsamples in SJLIFE, where treatment exposure was defined as any exposure to one or more curative agents for pediatric cancer previously associated with the specific phenotype.

In order to evaluate whether the observed replication frequencies were greater or less than expected for each of our phenotypes, we used a Poisson generalized estimating equations (GEE) regression approach with robust variance estimation^53^. We estimated the expected number of replications for each phenotype based on the assumption that each SNP replication may be treated as a Bernoulli random variable with a replication probability equal to its estimated replication power, and under Le Cam’s theorem^54^, the sum of independent Bernoulli variables that are not identically distributed approximately follows a Poisson distribution. The model assumed a log-link of the following form:

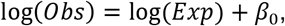

where *obs* and *Exp* were observed replications and the expected replication probability, respectively. The exponentiated *β*_0_ estimate served as the Replication Enrichment Ratio (RER), or the ratio of observed to expected replication frequencies.

### Ancillary analyses: Epigenetic and functional annotation enrichments by SNP replication state

We applied epigenetic/functional annotations using resources provided by Roadmap Epigenomics Mapping Consortium^18^ (REMC), Genotype-Tissue Expression Project^17^ (GTEx Analysis v7), Reactome^20^, and BIOS QTL^21^. We assessed the specificity of enhancer and promoter states for all SNPs with at least one replicated association in the SJLIFE main analysis using the REMC 15-chromatin state annotation data for 127 human cell types. For each cell type, we compared the frequency of enhancer/promoter state overlap in the set of SNPs with replicated associations (“replicated SNPs”) against the SNPs without replicated associations (“non-replicated SNPs”) in our SJLIFE main analysis. We evaluated nominal enrichment for these regulatory states using *P*-values obtained from 2-sided Fisher’s exact tests. Using GTEx, we counted the number of significant *cis*-eQTLs (SNPs within ±1 Mb of transcription start sites, FDR≤0.05) for replicated SNPs and non-replicated SNPs and used a 2-sided Fisher’s exact test to investigate enrichments in gene expressions among replicated SNPs for each of the 48 available cell-/tissue-types. Lastly, we compiled non-overlapping gene sets for replicated SNPs and non-replicated SNPs to conduct a biological pathway enrichment analysis with geneSCF v1.1^55^ and Reactome gene pathway ontologies. A gene was regarded as relevant to a SNP group if a SNP was located within the body of a RefSeq^19^ gene. For each biological pathway, the number of genes in our SNP groups with that ontology were compared to the number of genes with that ontology in all remaining genes in the genome. Top biological pathway enrichments were determined using FDR-adjusted *P*-values from 2-sided Fisher’s exact tests. Lastly, we used BIOS QTL^21^ to identify significant (FDR<0.05) *cis*-meQTLs linked to our 1,231 meta-GWAS SNPS and tested for enrichments/depletions of SNPs with ≥1 *cis-*meQTL among the replicated and non-replicated SNPs in our SJLIFE main analysis with two-sided Fisher’s exact tests.

### SNP-methylation and treatment-methylation associations

As a first step, we sought to validate significant (FDR<5%) *cis*-meQTLs reported in BIOS QTL in our sample of 236 SJLIFE participants with methylation and genotype data. For each established *cis*-meQTL available for testing in SJLIFE, we considered M-values (log_2_-transformed ratio of the methylated to unmethylated probe intensities) at quality-controlled CpG sites and tested associations between CpG M-values and SNP genotypes assuming an additive inheritance model using linear regression, adjusting for sex, age, and genetic ancestry. Since additional analyses to evaluate potential confounding by inter-individual differences in blood cell composition revealed no significant differences in cell type distributions across samples, no adjustment covariates for blood cell composition were considered. Established *cis-*meQTLs (i.e., reported in BIOS QTL with FDR<5%) were defined as validated in SJLIFE for associations with *P*<0.05 and the same direction of allelic effect.

We tested for enrichment of SJLIFE-validated *cis-*meQTLs among non-replicated SNPs with at least one significant *cis-*meQTL in BIOS QTL using a 2-sided Fisher’s exact test. We also identified *a priori* 48 “treatment-sensitive” meta-GWAS SNPs (without replications in our main analysis but were replicated in samples without treatment exposures) and 66 “treatment-insensitive” meta-GWAS SNPs (replicated in treatment-unexposed and treatment-exposed samples) and tested for enrichment of validated *cis-*meQTLs among treatment-sensitive SNPs. Finally, we examined directionally discordant SNP-methylation and treatment-methylation associations for CpGs linked to non-replicated SNPs (“non-replicated CpGs”) and CpGs linked to replicated SNPs (“replicated CpGs”) for the *cis-*meQTLs we validated in SJLIFE. Among the eight treatment types we considered (cranial, chest, abdominal, and pelvic radiotherapies; anthracycline, corticosteroid, cisplatin, and carboplatin chemotherapies), we limited our analysis to seven treatment types where >5% of the experimental sample was exposed. To ascertain the direction of SNP-CpG methylation associations for CpGs in SJLIFE-validated meQTLs with multiple associated SNPs without arbitrarily assigning a “best” SNP-CpG (i.e., smallest *P*-value), we used simple majority voting classification to assign the direction of the SNP-methylation association for such CpGs. For each treatment type, treatment dose associations with M-values at CpGs contributing to SJLIFE-validated *cis*-meQTLs were tested with linear regression, adjusting for age and sex. We compared the discordance between directions of SNP-methylation and treatment-methylation associations at each CpG for each of the seven treatment types among replicated and non-replicated CpGs using a two-sided Fisher’s exact test.

## DATA AVAILABILITY

The data used in this study may be accessed from the St. Jude Cloud (https://www.stjude.cloud/) under accession number SJC-DS-1002.

## ACKNOWLEDGEMENTS

This work was funded by the National Cancer Institute (U24 CA55727 GT Armstrong, principal investigator, U01 CA195547 MM Hudson/L Robison, principal investigators, CA 21765, C. Roberts, Principal Investigator and R01 CA216354), American Lebanese Syrian Associated Charities, and Alberta Machine Intelligence Institute.

## AUTHOR CONTRIBUTIONS

C.I., Y.Y. designed study concept and analytic methodologies. W.C., T.M.G., D.A.M., C.L.W. informed phenotype models specific to survivors. C.I., Y.Y., W.Q., N.Q., Z.W. performed analyses. Z.W., J.Z., W.C., K.K.N., C.L.W., Y.S., W.M., M.M.H., L.L.R., L.M.M., G.T.A. oversaw recruitment, sample collection, genotyping/sequencing, and data processing in SJLIFE and CCSS studies. Z.W., N.Q., J.Z. coordinated the generation and processing of methylome data. C.I., C.R.H., K.K.N., W.Q. managed phenotype, clinical data. C.I., N.Q., Z.W., Y.Y. drafted the paper. All authors critically revised and approved the final manuscript.

