## Supplemental Information for "Generalizability of “GWAS hits” in clinical populations: Lessons from childhood cancer survivors"

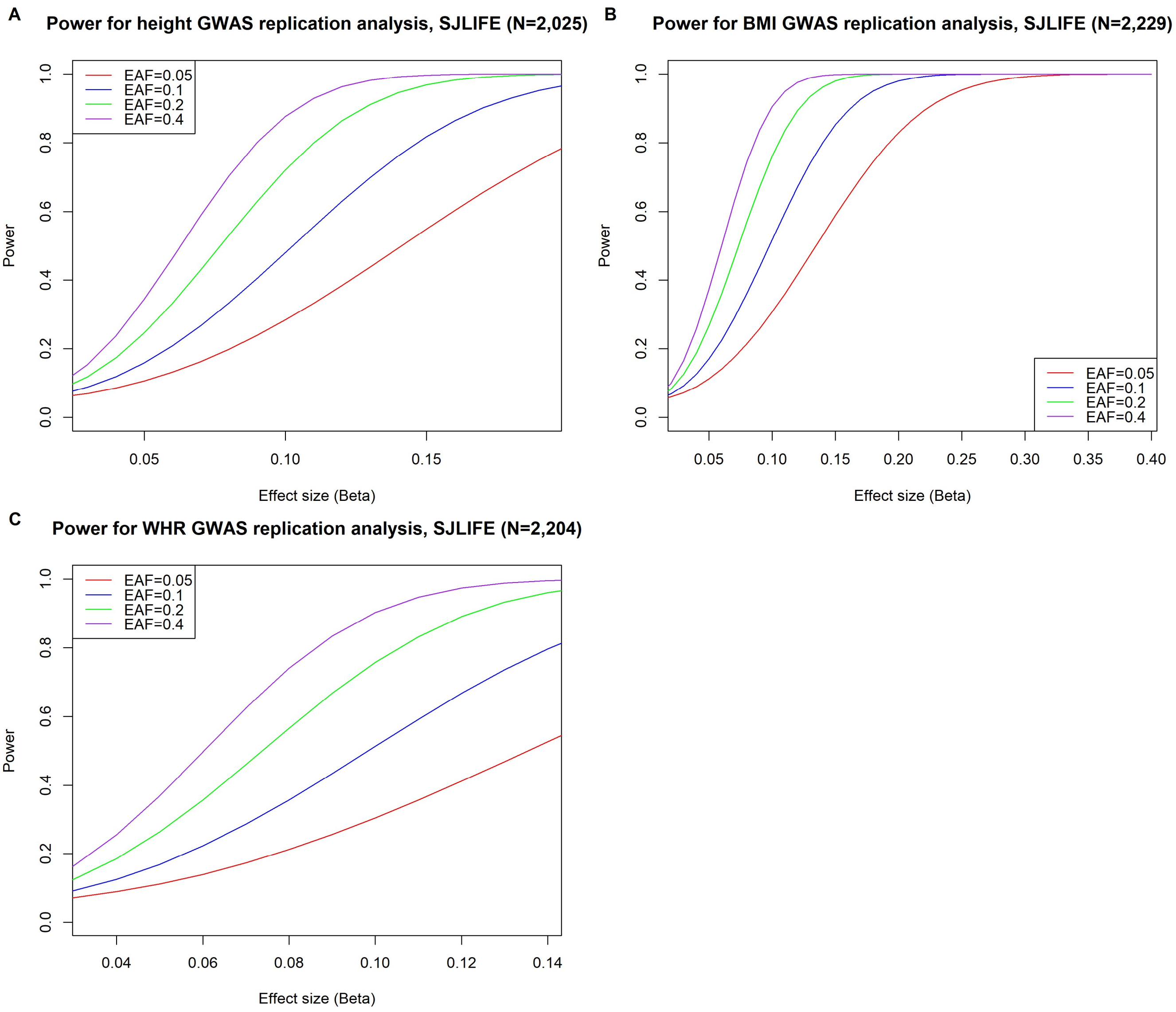


**Supplementary Figure 1**: Power analysis curves for anthropometric traits (A: height; B: body mass index [BMI]; and C: waist-to-hip ratio [WHR]) in the SJLIFE cohort for various effect allele frequencies (EAF, represented by colored curves) and effect sizes, assuming an additive genetic inheritance model and a 5% significance level.

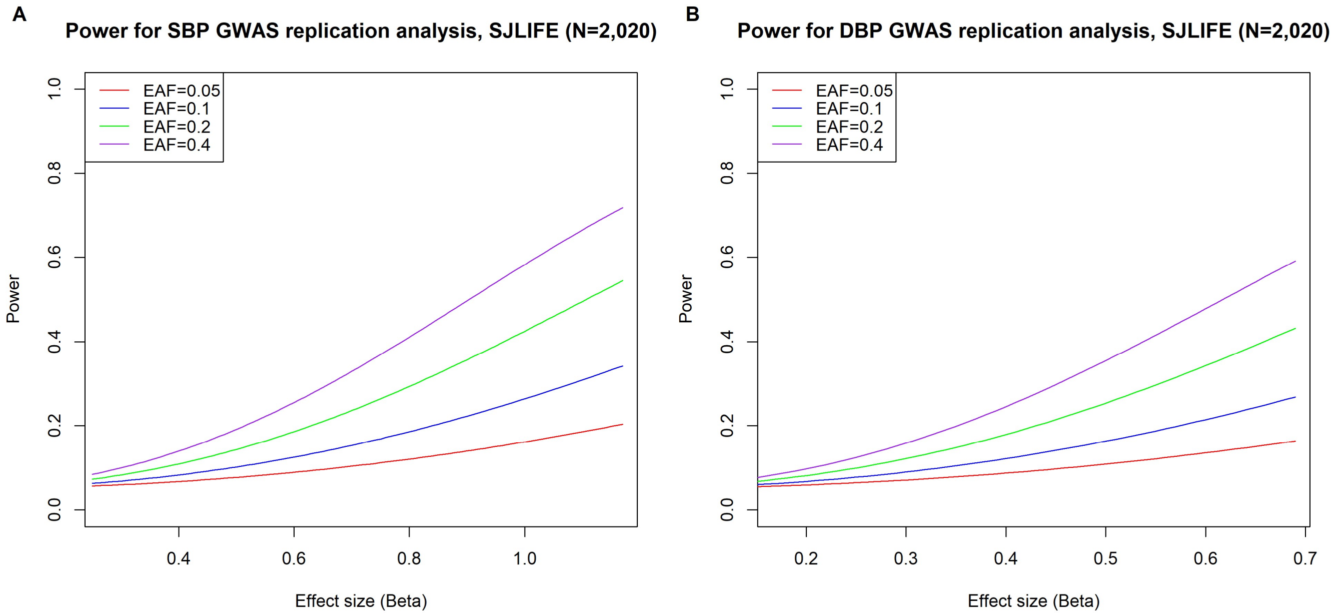


**Supplementary Figure 2**: Power analysis curves for blood pressure (A: systolic blood pressure [SBP]; B: diastolic blood pressure [DBP]) in the SJLIFE cohort for various effect allele frequencies (EAF, represented by colored curves) and effect sizes, assuming an additive genetic inheritance model and a 5% significance level.

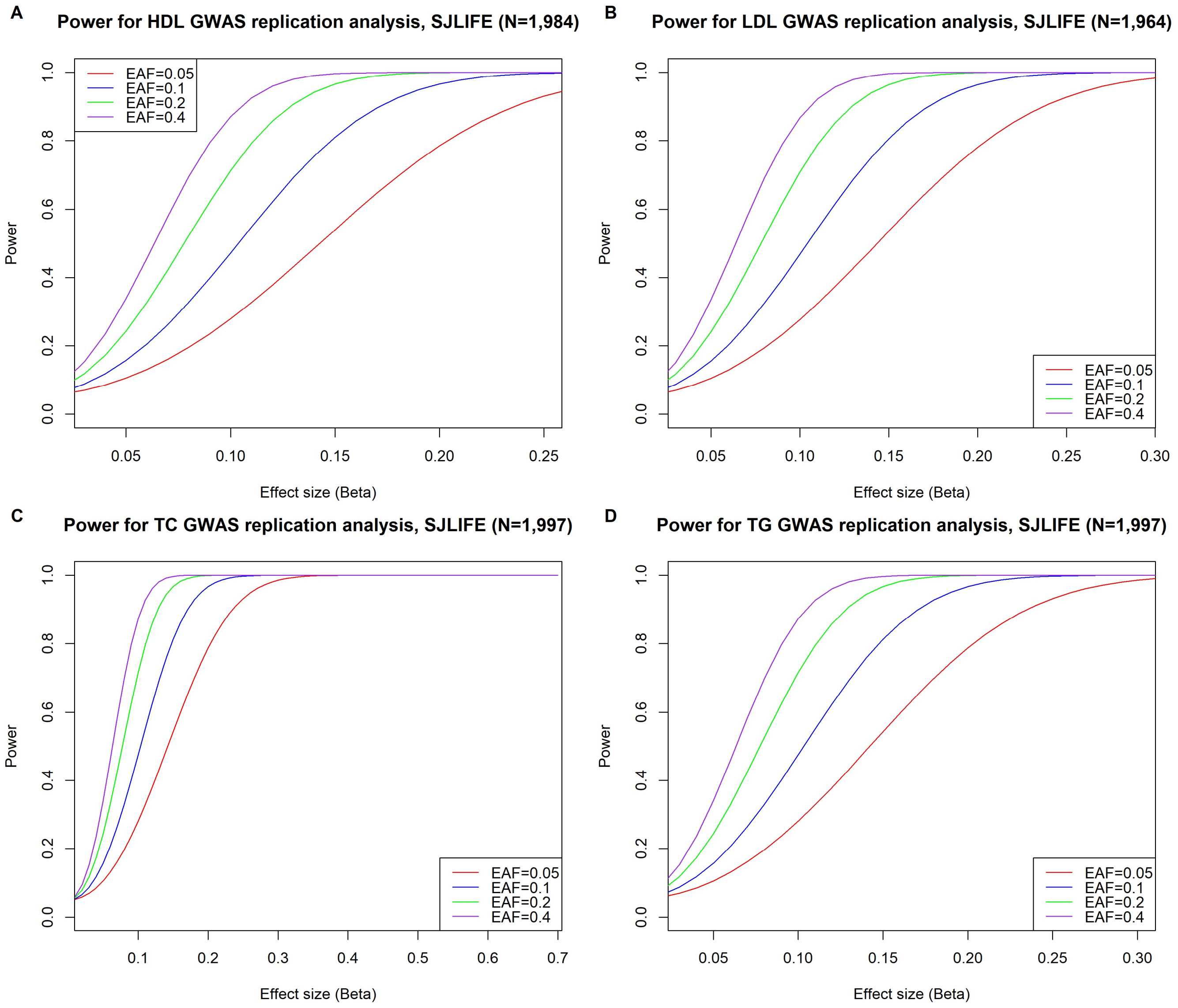


**Supplementary Figure 3**: Power analysis curves for blood lipid traits (A: high density lipoprotein [HDL]; B: low density lipoprotein [LDL]; C: total cholesterol [TC]; D: triglycerides [TG]) in the SJLIFE cohort for various effect allele frequencies (EAF, represented by colored curves) and effect sizes, assuming an additive genetic inheritance model and a 5% significance level.

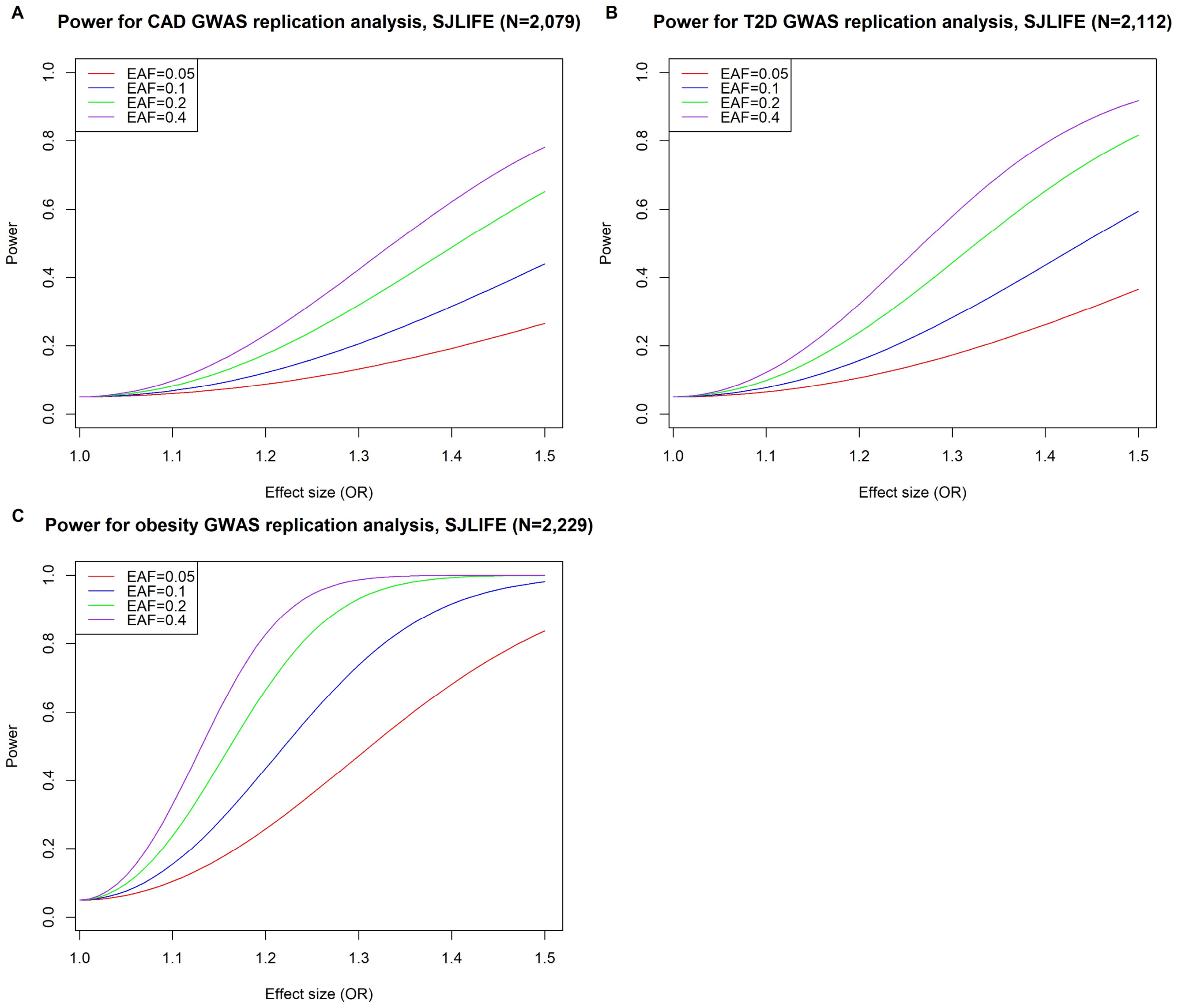


**Supplementary Figure 4**: Power analysis curves for cardiovascular and metabolic disease phenotypes (A: coronary artery disease [CAD]; B: type 2 diabetes [T2D]; C: obesity) in the SJLIFE cohort for various effect allele frequencies (EAF, represented by colored curves) and effect sizes, assuming an additive genetic inheritance model and a 5% significance level.

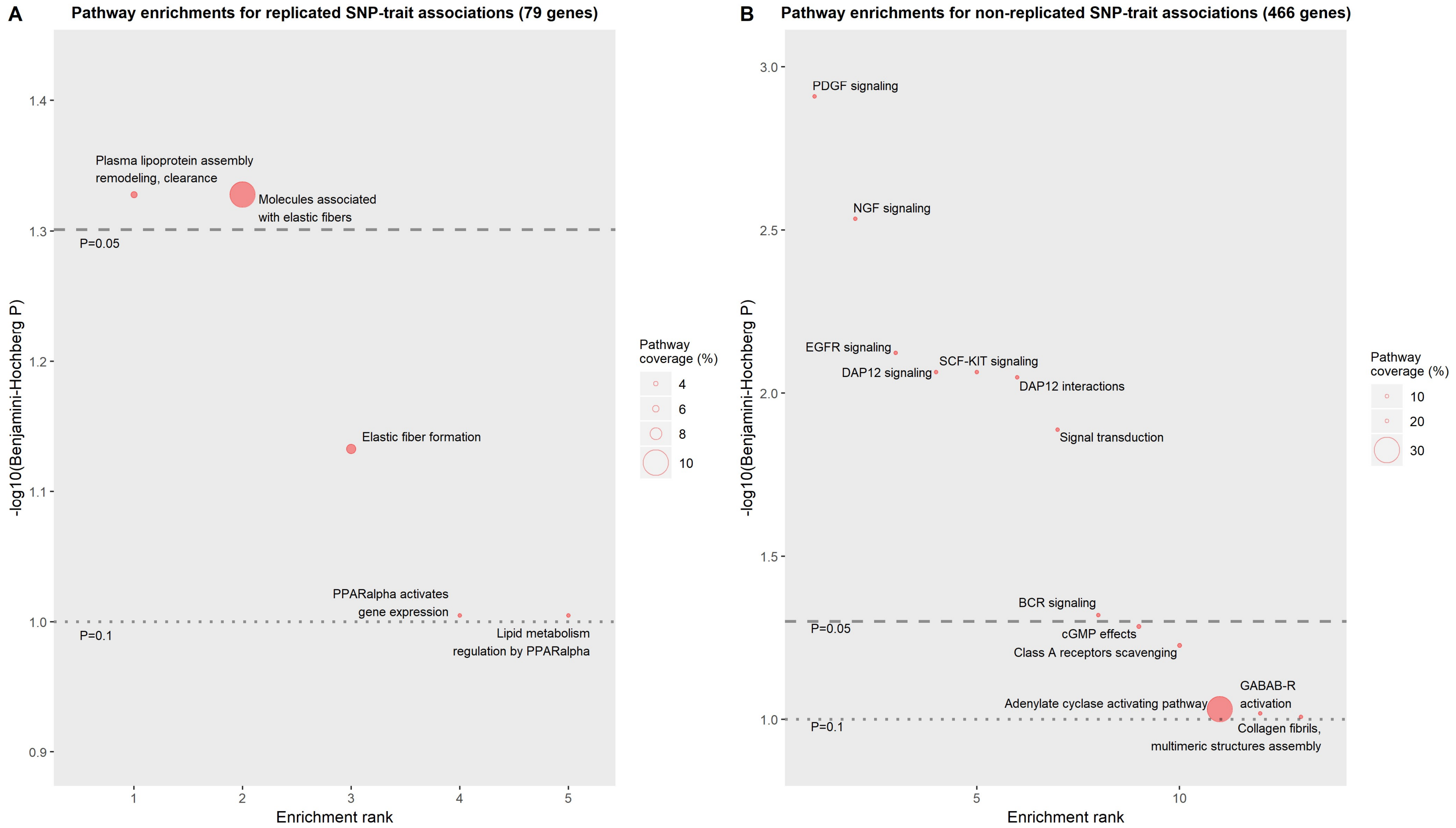


**Supplementary Figure 5**: Top Reactome pathway enrichment results for genes linked to replicated SNP-trait associations (A) and non-replicated SNP-trait associations (B). Top results (i.e., "enrichment rank") are defined by smaller FDR-adjusted (Benjamini-Hochberg) *P*-values. All results with FDR-adjusted *P*<0.1 are shown. Contributing genes for SNPs of interest were defined by whether SNPs of interest resided in RefSeq gene bodies. Plotted *P*-values are from 2-sided Fisher's exact tests comparing number of genes linked to the replicated/non-replicated SNP set with a given Reactome pathway ontology against the number of genes with the same ontology among all genes in the human genome. The circle sizes of plotted *P*-values correspond to the percent coverage of a given Reactome biological pathway by the genes with the same ontology linked to replicated/non-replicated SNP sets. Dashed and dotted lines correspond to FDRs of 5% and 10%, respectively.

**Supplementary Table 1**: Replicated SNP-phenotype associations in the SJLIFE cohort, adjusted for covariates used in reference meta-GWAS

| Phenotype | SNP | Chr | BP | EA | SJLIFE EAF | SJLIFE Beta/OR | SJLIFE P | GWAS-reported EAF | GWAS-reported Beta/OR | GWAS-reported P | Power |
|---|---|---|---|---|---|---|---|---|---|---|---|
| BMI | rs1558902 | 16 | 53769662 | A | 0.40 | 0.12 | 1.2E-04 | 0.42 | 0.39 | 5.0E-120 | 1.00 |
| BMI | rs9940128 | 16 | 53766842 | A | 0.42 | 0.11 | 1.5E-04 | 0.43 | 0.08 | 4.0E-23 | 0.75 |
| BMI | rs7138803 | 12 | 49853685 | A | 0.37 | 0.12 | 1.8E-04 | 0.38 | 0.12 | 2.0E-17 | 0.97 |
| BMI | rs62033400 | 16 | 53777876 | G | 0.39 | 0.11 | 2.6E-04 | 0.42 | 0.08 | 2.0E-14 | 0.74 |
| BMI | rs9939609 | 16 | 53786615 | A | 0.40 | 0.11 | 3.4E-04 | 0.41 | 0.33 | 4.0E-51 | 1.00 |
| BMI | rs17782313 | 18 | 60183864 | C | 0.24 | 0.13 | 3.5E-04 | 0.21 | 0.20 | 5.0E-18 | 1.00 |
| BMI | rs6567160 | 18 | 60161902 | C | 0.23 | 0.13 | 3.8E-04 | 0.24 | 0.09 | 8.0E-19 | 0.72 |
| BMI | rs571312 | 18 | 60172536 | A | 0.24 | 0.12 | 4.3E-04 | 0.24 | 0.23 | 6.0E-42 | 1.00 |
| BMI | rs7202116 | 16 | 53787703 | G | 0.40 | 0.11 | 4.4E-04 | 0.40 | 0.04 | 2.0E-10 | 0.26 |
| BMI | rs12286929 | 11 | 115151684 | G | 0.54 | 0.09 | 2.3E-03 | 0.52 | 0.02 | 1.0E-12 | 0.10 |
| BMI | rs977747 | 1 | 47219005 | T | 0.41 | 0.09 | 3.9E-03 | 0.40 | 0.02 | 2.0E-08 | 0.10 |
| BMI | rs2075650 | 19 | 44892362 | A | 0.86 | 0.11 | 0.01 | 0.85 | 0.03 | 1.0E-08 | 0.11 |
| BMI | rs13191362 | 6 | 162612318 | A | 0.88 | 0.12 | 0.01 | 0.88 | 0.03 | 1.0E-09 | 0.10 |
| BMI | rs12940622 | 17 | 80641771 | G | 0.56 | 0.07 | 0.02 | 0.58 | 0.02 | 2.0E-09 | 0.10 |
| BMI | rs7234864 | 18 | 60067625 | T | 0.27 | 0.07 | 0.04 | 0.26 | 0.08 | 4.0E-17 | 0.65 |
| CAD | rs1412444 | 10 | 89243170 | T | 0.34 | 1.51 | 0.01 | 0.37 | 1.07 | 5.0E-12 | 0.07 |
| CAD | rs2246942 | 10 | 89245129 | G | 0.35 | 1.46 | 0.01 | 0.35 | 1.08 | 4.0E-16 | 0.08 |
| CAD | rs1332329 | 10 | 89243662 | C | 0.35 | 1.45 | 0.02 | 0.36 | 1.08 | 3.0E-13 | 0.08 |
| CAD | rs8108632 | 19 | 41348629 | T | 0.43 | 1.40 | 0.03 | 0.48 | 1.05 | 4.0E-08 | 0.06 |
| DBP | rs62524579 | 8 | 142979538 | A | 0.54 | -0.80 | 0.01 | 0.53 | -0.18 | 4.0E-09 | 0.09 |
| DBP | rs3184504 | 12 | 111446804 | T | 0.48 | 0.73 | 0.01 | 0.48 | 0.48 | 3.0E-14 | 0.34 |
| DBP | rs653178 | 12 | 111569952 | T | 0.51 | -0.70 | 0.01 | 0.53 | -0.46 | 3.0E-18 | 0.32 |
| HDL | rs173539 | 16 | 56954132 | T | 0.33 | 0.23 | 2.0E-11 | 0.32 | 0.25 | 4.0E-75 | 1.00 |
| HDL | rs3764261 | 16 | 56959412 | A | 0.33 | 0.23 | 2.2E-11 | 0.32 | 0.24 | 0.0E+00 | 1.00 |
| HDL | rs12678919 | 8 | 19986711 | G | 0.11 | 0.18 | 4.2E-04 | 0.10 | 0.23 | 2.0E-34 | 1.00 |
| HDL | rs1800961 | 20 | 44413724 | T | 0.03 | -0.28 | 1.9E-03 | 0.03 | -0.19 | 8.0E-10 | 0.53 |
| HDL | rs10468017 | 15 | 58386313 | T | 0.31 | 0.11 | 2.3E-03 | 0.30 | 0.10 | 8.0E-23 | 0.83 |
| HDL | rs16942887 | 16 | 67894139 | A | 0.13 | 0.14 | 3.1E-03 | 0.14 | 0.08 | 8.0E-54 | 0.40 |
| HDL | rs2271293 | 16 | 67868167 | A | 0.13 | 0.13 | 4.8E-03 | 0.11 | 0.07 | 9.0E-13 | 0.32 |
| HDL | rs4846914 | 1 | 230159944 | G | 0.41 | -0.09 | 5.0E-03 | 0.40 | -0.05 | 4.0E-08 | 0.34 |
| HDL | rs12328675 | 2 | 164684290 | C | 0.12 | 0.13 | 0.01 | 0.13 | 0.05 | 2.0E-15 | 0.13 |
| HDL | rs9987289 | 8 | 9325848 | A | 0.09 | -0.15 | 0.01 | 0.10 | -0.08 | 2.0E-41 | 0.30 |
| HDL | rs1883025 | 9 | 104902020 | T | 0.27 | -0.08 | 0.02 | 0.26 | -0.08 | 1.0E-09 | 0.61 |
| HDL | rs386000 | 19 | 54288907 | C | 0.21 | 0.08 | 0.03 | 0.26 | 0.05 | 3.0E-23 | 0.25 |
| HDL | rs7255436 | 19 | 8368312 | C | 0.49 | -0.07 | 0.03 | 0.47 | -0.03 | 2.0E-08 | 0.16 |
| HDL | rs7679 | 20 | 45947863 | C | 0.19 | -0.09 | 0.04 | 0.19 | -0.07 | 4.0E-09 | 0.41 |
| HDL | rs6065906 | 20 | 45925376 | C | 0.19 | -0.08 | 0.04 | 0.19 | -0.06 | 5.0E-40 | 0.32 |
| HDL | rs2602836 | 4 | 99093654 | A | 0.41 | 0.07 | 0.04 | 0.44 | 0.02 | 5.0E-08 | 0.10 |
| HDL | rs4983559 | 14 | 104810872 | G | 0.39 | 0.06 | 0.04 | 0.40 | 0.02 | 1.0E-08 | 0.09 |
| HDL | rs1532085 | 15 | 58391167 | A | 0.40 | 0.07 | 0.04 | 0.40 | 0.11 | 1.0E-188 | 0.93 |
| Height | rs2961830 | 5 | 51158898 | A | 0.35 | 0.11 | 1.0E-03 | 0.35 | 0.02 | 2.4E-10 | 0.09 |
| Height | rs1812175 | 4 | 144653692 | A | 0.17 | -0.14 | 1.2E-03 | 0.16 | -0.08 | 2.0E-86 | 0.48 |
| Height | rs7689420 | 4 | 144647200 | T | 0.17 | -0.13 | 1.8E-03 | 0.16 | -0.07 | 6.0E-51 | 0.39 |
| Height | rs7763064 | 6 | 142476152 | A | 0.32 | -0.10 | 2.1E-03 | 0.29 | -0.05 | 1.0E-33 | 0.32 |
| Height | rs1550162 | 8 | 116551294 | A | 0.73 | -0.10 | 4.0E-03 | 0.71 | -0.02 | 1.3E-13 | 0.09 |
| Height | rs17511102 | 2 | 37733470 | A | 0.92 | -0.17 | 4.0E-03 | 0.91 | -0.06 | 2.0E-18 | 0.18 |
| Height | rs3748069 | 6 | 142446496 | A | 0.69 | 0.10 | 4.1E-03 | 0.74 | 0.07 | 5.0E-14 | 0.42 |
| Height | rs4896582 | 6 | 142382740 | A | 0.33 | -0.09 | 4.4E-03 | 0.27 | -0.05 | 2.0E-18 | 0.32 |
| Height | rs1036821 | 8 | 134638240 | A | 0.32 | -0.09 | 0.01 | 0.30 | -0.04 | 1.0E-30 | 0.22 |
| Height | rs11830103 | 12 | 123338999 | A | 0.79 | -0.10 | 0.01 | 0.78 | -0.04 | 4.0E-15 | 0.18 |
| Height | rs3807931 | 7 | 20342051 | A | 0.46 | 0.08 | 0.01 | 0.45 | 0.03 | 1.2E-19 | 0.16 |
| Height | rs2834442 | 21 | 34318486 | A | 0.65 | 0.09 | 0.01 | 0.65 | 0.03 | 5.0E-12 | 0.15 |
| Height | rs2305833 | 2 | 218440681 | C | 0.57 | 0.08 | 0.01 | 0.58 | 0.03 | 3.8E-21 | 0.16 |
| Height | rs7980687 | 12 | 123338164 | A | 0.20 | 0.10 | 0.01 | 0.21 | 0.04 | 1.0E-26 | 0.17 |
| Height | rs2154319 | 1 | 41280098 | T | 0.78 | -0.10 | 0.01 | 0.75 | -0.03 | 2.0E-12 | 0.12 |
| Height | rs3118905 | 13 | 50531198 | A | 0.28 | -0.09 | 0.01 | 0.28 | -0.06 | 1.0E-69 | 0.40 |
| Height | rs10997979 | 10 | 68177435 | A | 0.51 | -0.08 | 0.01 | 0.50 | -0.02 | 4.0E-13 | 0.10 |
| Height | rs9409082 | 9 | 106138768 | T | 0.24 | -0.09 | 0.01 | 0.24 | -0.03 | 8.8E-15 | 0.13 |
| Height | rs6440003 | 3 | 141375367 | A | 0.43 | 0.08 | 0.02 | 0.44 | 0.07 | 2.0E-24 | 0.60 |
| Height | rs6763931 | 3 | 141383991 | A | 0.43 | 0.07 | 0.02 | 0.45 | 0.07 | 1.0E-27 | 0.60 |
| Height | rs3020418 | 6 | 152024027 | A | 0.28 | 0.08 | 0.02 | 0.30 | 0.03 | 8.0E-24 | 0.14 |
| Height | rs3764419 | 17 | 30837005 | A | 0.39 | -0.07 | 0.02 | 0.39 | -0.04 | 2.0E-21 | 0.24 |
| Height | rs12470505 | 2 | 219043647 | T | 0.89 | 0.12 | 0.02 | 0.90 | 0.05 | 6.2E-22 | 0.17 |
| Height | rs16994718 | 4 | 38686741 | T | 0.16 | -0.10 | 0.02 | 0.15 | -0.03 | 1.6E-09 | 0.08 |
| Height | rs4256170 | 3 | 51154695 | A | 0.03 | 0.23 | 0.02 | 0.01 | 0.19 | 1.0E-11 | 0.54 |
| Height | rs2284746 | 1 | 16980180 | C | 0.48 | -0.07 | 0.02 | 0.48 | -0.04 | 4.0E-29 | 0.25 |
| Height | rs2237886 | 11 | 2789501 | T | 0.11 | 0.12 | 0.02 | 0.11 | 0.05 | 2.0E-13 | 0.17 |
| Height | rs12615742 | 2 | 37768584 | T | 0.48 | 0.07 | 0.02 | 0.48 | 0.03 | 2.7E-13 | 0.16 |
| Height | rs4965598 | 15 | 100219409 | T | 0.68 | -0.08 | 0.02 | 0.68 | -0.03 | 4.0E-13 | 0.14 |
| Height | rs3760318 | 17 | 30920697 | A | 0.38 | -0.07 | 0.02 | 0.37 | -0.04 | 3.0E-41 | 0.24 |

| Phenotype | SNP | Chr | BP | EA | SJLIFE EAF | SJLIFE Beta/OR | SJLIFE P | GWAS-reported EAF | GWAS-reported Beta/OR | GWAS-reported P | Power |
|---|---|---|---|---|---|---|---|---|---|---|---|
| Height | rs724016 | 3 | 141386728 | G | 0.43 | 0.07 | 0.02 | 0.48 | 0.05 | 8.0E-22 | 0.35 |
| Height | rs10010325 | 4 | 105185196 | A | 0.49 | 0.07 | 0.02 | 0.49 | 0.02 | 4.0E-11 | 0.10 |
| Height | rs10748128 | 12 | 69433878 | T | 0.35 | 0.07 | 0.03 | 0.35 | 0.04 | 2.0E-20 | 0.23 |
| Height | rs7926971 | 11 | 12676493 | A | 0.54 | -0.07 | 0.03 | 0.55 | -0.02 | 4.0E-10 | 0.10 |
| Height | rs9880211 | 3 | 136388707 | A | 0.24 | -0.08 | 0.03 | 0.25 | -0.03 | 2.0E-18 | 0.13 |
| Height | rs4369779 | 18 | 23155444 | T | 0.21 | -0.08 | 0.03 | 0.21 | -0.06 | 2.0E-53 | 0.34 |
| Height | rs2145272 | 20 | 6645571 | A | 0.64 | -0.07 | 0.03 | 0.65 | -0.04 | 2.0E-24 | 0.23 |
| Height | rs6485978 | 11 | 12656868 | T | 0.54 | -0.07 | 0.03 | 0.54 | -0.02 | 1.0E-15 | 0.10 |
| Height | rs13177718 | 5 | 108777643 | T | 0.08 | -0.13 | 0.03 | 0.08 | -0.04 | 3.0E-13 | 0.11 |
| Height | rs6724465 | 2 | 219079124 | A | 0.10 | -0.11 | 0.03 | 0.10 | -0.06 | 2.0E-08 | 0.21 |
| Height | rs2079795 | 17 | 61419288 | T | 0.35 | 0.07 | 0.03 | 0.33 | 0.05 | 2.0E-46 | 0.23 |
| Height | rs7727731 | 5 | 65378619 | T | 0.13 | 0.10 | 0.03 | 0.11 | 0.03 | 1.3E-11 | 0.10 |
| Height | rs7759938 | 6 | 104931079 | T | 0.67 | -0.07 | 0.03 | 0.68 | -0.05 | 8.0E-31 | 0.22 |
| Height | rs17410035 | 5 | 31541035 | T | 0.33 | 0.07 | 0.03 | 0.33 | 0.02 | 9.0E-10 | 0.09 |
| Height | rs1013209 | 8 | 24258791 | T | 0.23 | -0.08 | 0.03 | 0.25 | -0.03 | 2.0E-09 | 0.08 |
| Height | rs1809889 | 12 | 124316680 | T | 0.26 | 0.08 | 0.03 | 0.29 | 0.03 | 4.0E-21 | 0.13 |
| Height | rs1265097 | 6 | 31138682 | A | 0.11 | -0.11 | 0.03 | 0.12 | -0.06 | 2.0E-32 | 0.22 |
| Height | rs1753637 | 13 | 50510037 | T | 0.30 | 0.07 | 0.03 | 0.31 | 0.04 | 1.1E-29 | 0.21 |
| Height | rs6845999 | 4 | 144644674 | T | 0.43 | 0.07 | 0.04 | 0.44 | 0.05 | 5.0E-67 | 0.35 |
| Height | rs8073371 | 17 | 48018910 | T | 0.21 | -0.08 | 0.04 | 0.22 | -0.02 | 3.8E-12 | 0.08 |
| Height | rs10790381 | 11 | 120386786 | A | 0.85 | 0.09 | 0.04 | 0.82 | 0.03 | 2.0E-12 | 0.10 |
| Height | rs4743034 | 9 | 106870072 | A | 0.23 | 0.08 | 0.04 | 0.23 | 0.05 | 2.0E-08 | 0.27 |
| Height | rs314263 | 6 | 104944870 | T | 0.67 | -0.07 | 0.04 | 0.68 | -0.04 | 1.0E-42 | 0.22 |
| Height | rs1341278 | 6 | 80329204 | T | 0.94 | -0.14 | 0.04 | 0.94 | -0.06 | 5.8E-18 | 0.15 |
| Height | rs9844666 | 3 | 136255374 | A | 0.23 | -0.07 | 0.04 | 0.25 | -0.02 | 4.0E-09 | 0.08 |
| Height | rs1046934 | 1 | 184054395 | A | 0.66 | -0.07 | 0.04 | 0.64 | -0.04 | 2.0E-31 | 0.23 |
| Height | rs2145998 | 10 | 79361940 | A | 0.48 | -0.06 | 0.04 | 0.49 | -0.03 | 4.0E-13 | 0.16 |
| Height | rs17369123 | 1 | 172386701 | T | 0.18 | 0.08 | 0.04 | 0.19 | 0.03 | 1.4E-16 | 0.11 |
| Height | rs4735677 | 8 | 77235955 | A | 0.72 | -0.07 | 0.04 | 0.72 | -0.04 | 6.0E-30 | 0.21 |
| Height | rs4800148 | 18 | 23144364 | A | 0.78 | 0.08 | 0.04 | 0.79 | 0.06 | 4.0E-09 | 0.35 |
| Height | rs7853377 | 9 | 83937290 | A | 0.79 | -0.08 | 0.04 | 0.77 | -0.02 | 5.0E-08 | 0.08 |
| Height | rs4800452 | 18 | 23147647 | T | 0.78 | 0.07 | 0.05 | 0.79 | 0.05 | 4.0E-30 | 0.26 |
| Height | rs9863706 | 3 | 72388262 | T | 0.21 | -0.08 | 0.05 | 0.22 | -0.03 | 4.0E-13 | 0.12 |
| Height | rs7846385 | 8 | 77247943 | C | 0.28 | 0.07 | 0.05 | 0.27 | 0.05 | 5.0E-08 | 0.30 |
| Height | rs2341459 | 2 | 44541063 | T | 0.26 | 0.07 | 0.05 | 0.27 | 0.03 | 8.0E-10 | 0.09 |
| Height | rs13088462 | 3 | 51034282 | T | 0.95 | -0.14 | 0.05 | 0.94 | -0.06 | 8.0E-18 | 0.13 |
| Height | rs3818416 | 13 | 77900333 | A | 0.22 | -0.08 | 0.05 | 0.22 | -0.02 | 1.7E-09 | 0.08 |
| Height | rs3811958 | 5 | 32771937 | A | 0.74 | -0.07 | 0.05 | 0.74 | -0.03 | 2.7E-16 | 0.13 |
| LDL | rs12740374 | 1 | 109274968 | T | 0.23 | -0.17 | 3.6E-06 | 0.21 | -0.23 | 2.0E-42 | 1.00 |
| LDL | rs4420638 | 19 | 44919689 | G | 0.18 | 0.19 | 3.8E-06 | 0.16 | 0.29 | 4.0E-27 | 1.00 |
| LDL | rs629301 | 1 | 109275684 | G | 0.23 | -0.17 | 4.2E-06 | 0.24 | -0.17 | 5.0E-241 | 0.99 |
| LDL | rs6511720 | 19 | 11091630 | T | 0.12 | -0.21 | 3.8E-05 | 0.10 | -0.26 | 2.0E-26 | 1.00 |
| LDL | rs11206510 | 1 | 55030366 | C | 0.18 | -0.16 | 2.1E-04 | 0.19 | -0.09 | 4.0E-08 | 0.58 |
| LDL | rs10401969 | 19 | 19296909 | C | 0.07 | -0.18 | 2.9E-03 | 0.09 | -0.12 | 3.0E-54 | 0.48 |
| LDL | rs635634 | 9 | 133279427 | T | 0.19 | 0.12 | 3.5E-03 | 0.21 | 0.08 | 2.0E-41 | 0.50 |
| LDL | rs6882076 | 5 | 156963286 | T | 0.36 | -0.09 | 3.7E-03 | 0.36 | -0.05 | 3.0E-31 | 0.32 |
| LDL | rs1501908 | 5 | 156971158 | G | 0.36 | -0.09 | 0.01 | 0.37 | -0.07 | 1.0E-11 | 0.56 |
| LDL | rs174546 | 11 | 61802358 | T | 0.33 | -0.08 | 0.01 | 0.36 | -0.05 | 2.0E-39 | 0.31 |
| LDL | rs2954029 | 8 | 125478730 | T | 0.47 | -0.08 | 0.01 | 0.47 | -0.06 | 2.0E-50 | 0.47 |
| LDL | rs1169288 | 12 | 120978847 | C | 0.33 | 0.08 | 0.02 | 0.34 | 0.04 | 6.0E-21 | 0.22 |
| LDL | rs1564348 | 6 | 160157828 | C | 0.16 | 0.10 | 0.03 | 0.18 | 0.05 | 3.0E-21 | 0.21 |
| LDL | rs515135 | 2 | 21063185 | T | 0.19 | -0.09 | 0.03 | 0.20 | -0.16 | 5.0E-29 | 0.98 |
| LDL | rs2479409 | 1 | 55038977 | G | 0.35 | 0.07 | 0.04 | 0.32 | 0.06 | 3.0E-50 | 0.43 |
| Obesity | rs538656 | 18 | 60183189 | T | 0.24 | 1.29 | 5.2E-04 | 0.24 | 1.15 | 2.0E-36 | 0.50 |
| Obesity | rs10871777 | 18 | 60184530 | G | 0.24 | 1.28 | 7.9E-04 | 0.24 | 1.10 | 2.0E-27 | 0.26 |
| Obesity | rs11152213 | 18 | 60185715 | C | 0.24 | 1.27 | 1.0E-03 | 0.24 | 1.19 | 3.0E-22 | 0.68 |
| Obesity | rs1558902 | 16 | 53769662 | A | 0.40 | 1.21 | 3.1E-03 | 0.41 | 1.14 | 2.0E-81 | 0.55 |
| Obesity | rs1421085 | 16 | 53767042 | C | 0.40 | 1.21 | 3.1E-03 | 0.41 | 1.45 | 6.0E-39 | 1.00 |
| Obesity | rs7185735 | 16 | 53788739 | G | 0.40 | 1.21 | 3.3E-03 | 0.40 | 1.33 | 1.0E-79 | 1.00 |
| Obesity | rs8043757 | 16 | 53779538 | T | 0.40 | 1.20 | 4.3E-03 | 0.40 | 1.23 | 5.0E-110 | 0.91 |
| Obesity | rs7138803 | 12 | 49853685 | A | 0.37 | 1.14 | 0.04 | 0.38 | 1.14 | 1.0E-16 | 0.54 |
| SBP | rs1458038 | 4 | 80243569 | T | 0.29 | 1.13 | 0.01 | 0.29 | 0.71 | 2.0E-23 | 0.30 |
| SBP | rs1378942 | 15 | 74785026 | C | 0.34 | 0.99 | 0.02 | 0.35 | 0.61 | 6.0E-23 | 0.25 |
| SBP | rs4590817 | 10 | 61707795 | G | 0.84 | 1.24 | 0.02 | 0.84 | 0.65 | 4.0E-12 | 0.18 |
| SBP | rs2681492 | 12 | 89619312 | T | 0.83 | 1.15 | 0.03 | 0.80 | 0.85 | 4.0E-11 | 0.29 |
| SBP | rs1011018 | 7 | 139763465 | A | 0.19 | -1.07 | 0.04 | 0.20 | -0.33 | 2.0E-08 | 0.09 |
| SBP | rs17249754 | 12 | 89666809 | G | 0.83 | 1.11 | 0.04 | 0.84 | 0.93 | 2.0E-18 | 0.34 |
| T2D | rs11123406 | 2 | 111192964 | C | 0.64 | 0.66 | 1.5E-03 | 0.64 | 0.96 | 8.6E-09 | 0.06 |
| T2D | rs2925979 | 16 | 81501185 | T | 0.31 | 1.50 | 1.9E-03 | 0.30 | 1.07 | 2.0E-09 | 0.08 |
| T2D | rs4689388 | 4 | 6268329 | A | 0.58 | 1.49 | 3.4E-03 | 0.62 | 1.08 | 1.3E-15 | 0.10 |
| T2D | rs10923931 | 1 | 119975336 | T | 0.10 | 1.65 | 0.01 | 0.11 | 1.13 | 4.0E-08 | 0.10 |
| T2D | rs576674 | 13 | 32980164 | A | 0.83 | 0.66 | 0.01 | 0.84 | 0.93 | 9.3E-13 | 0.08 |

| Phenotype | SNP | Chr | BP | EA | SJLIFE EAF | SJLIFE Beta/OR | SJLIFE P | GWAS-reported EAF | GWAS-reported Beta/OR | GWAS-reported P | Power |
|---|---|---|---|---|---|---|---|---|---|---|---|
| T2D | rs864745 | 7 | 28140937 | T | 0.50 | 1.38 | 0.01 | 0.50 | 1.10 | 5.0E-14 | 0.12 |
| T2D | rs1470579 | 3 | 185811292 | C | 0.30 | 1.38 | 0.01 | 0.30 | 1.14 | 1.4E-45 | 0.17 |
| T2D | rs3923113 | 2 | 164645339 | C | 0.35 | 0.72 | 0.02 | 0.41 | 0.93 | 1.3E-13 | 0.09 |
| T2D | rs3794991 | 19 | 19499787 | T | 0.08 | 1.61 | 0.02 | 0.08 | 1.08 | 4.1E-09 | 0.06 |
| T2D | rs2943641 | 2 | 226229029 | C | 0.64 | 1.34 | 0.03 | 0.63 | 1.19 | 9.0E-12 | 0.28 |
| T2D | rs2943640 | 2 | 226228869 | C | 0.65 | 1.34 | 0.04 | 0.64 | 1.08 | 5.1E-18 | 0.09 |
| T2D | rs12681990 | 8 | 37001668 | C | 0.17 | 1.38 | 0.04 | 0.15 | 1.05 | 3.1E-09 | 0.06 |
| T2D | rs340874 | 1 | 213985913 | C | 0.56 | 1.29 | 0.05 | 0.53 | 1.05 | 6.0E-10 | 0.07 |
| TC | rs4420638 | 19 | 44919689 | G | 0.18 | 0.18 | 2.0E-05 | 0.19 | 0.20 | 1.0E-149 | 1.00 |
| TC | rs1260326 | 2 | 27508073 | T | 0.40 | 0.13 | 4.0E-05 | 0.39 | 0.05 | 3.0E-42 | 0.34 |
| TC | rs629301 | 1 | 109275684 | G | 0.23 | -0.15 | 8.8E-05 | 0.24 | -0.13 | 2.0E-170 | 0.93 |
| TC | rs6511720 | 19 | 11091630 | T | 0.12 | -0.19 | 1.1E-04 | 0.12 | -0.19 | 5.0E-202 | 0.96 |
| TC | rs10401969 | 19 | 19296909 | C | 0.07 | -0.23 | 1.2E-04 | 0.09 | -0.14 | 4.0E-77 | 0.62 |
| TC | rs635634 | 9 | 133279427 | T | 0.19 | 0.12 | 2.8E-03 | 0.21 | 0.07 | 3.0E-35 | 0.41 |
| TC | rs1564348 | 6 | 160157828 | C | 0.16 | 0.12 | 0.01 | 0.18 | 0.05 | 3.0E-23 | 0.21 |
| TC | rs2954029 | 8 | 125478730 | T | 0.47 | -0.08 | 0.01 | 0.47 | -0.06 | 2.0E-65 | 0.47 |
| TC | rs6882076 | 5 | 156963286 | T | 0.36 | -0.08 | 0.02 | 0.36 | -0.05 | 5.0E-41 | 0.33 |
| TC | rs174546 | 11 | 61802358 | T | 0.33 | -0.08 | 0.02 | 0.36 | -0.05 | 3.0E-37 | 0.32 |
| TC | rs964184 | 11 | 116778201 | C | 0.86 | -0.10 | 0.03 | 0.84 | -0.12 | 3.0E-55 | 0.75 |
| TC | rs7570971 | 2 | 135080336 | A | 0.34 | 0.07 | 0.04 | 0.35 | 0.03 | 1.0E-13 | 0.15 |
| TC | rs1997243 | 7 | 1044141 | G | 0.15 | 0.09 | 0.04 | 0.16 | 0.03 | 3.0E-10 | 0.10 |
| TC | rs2072183 | 7 | 44539581 | C | 0.22 | 0.08 | 0.04 | 0.29 | 0.04 | 4.0E-15 | 0.18 |
| TG | rs12678919 | 8 | 19986711 | G | 0.11 | -0.31 | 1.2E-09 | 0.10 | -0.25 | 2.0E-41 | 1.00 |
| TG | rs964184 | 11 | 116778201 | G | 0.14 | 0.24 | 3.8E-07 | 0.14 | 0.30 | 4.0E-62 | 1.00 |
| TG | rs1260326 | 2 | 27508073 | T | 0.40 | 0.14 | 1.6E-05 | 0.45 | 0.12 | 2.0E-31 | 0.96 |
| TG | rs7557067 | 2 | 20985339 | G | 0.23 | -0.11 | 4.5E-03 | 0.22 | -0.08 | 9.0E-12 | 0.57 |
| TG | rs10401969 | 19 | 19296909 | C | 0.07 | -0.17 | 0.01 | 0.09 | -0.12 | 1.0E-69 | 0.49 |
| TG | rs17145738 | 7 | 73568544 | T | 0.14 | -0.13 | 0.01 | 0.13 | -0.12 | 9.0E-99 | 0.75 |
| TG | rs714052 | 7 | 73450539 | G | 0.14 | -0.12 | 0.01 | 0.12 | -0.16 | 3.0E-15 | 0.94 |
| TG | rs2954029 | 8 | 125478730 | T | 0.47 | -0.08 | 0.01 | 0.44 | -0.11 | 3.0E-19 | 0.94 |
| TG | rs2929282 | 15 | 43953733 | T | 0.05 | 0.18 | 0.02 | 0.07 | 0.07 | 2.0E-09 | 0.16 |
| TG | rs4846914 | 1 | 230159944 | G | 0.41 | 0.07 | 0.03 | 0.41 | 0.04 | 7.0E-31 | 0.24 |
| TG | rs7679 | 20 | 45947863 | C | 0.19 | 0.08 | 0.04 | 0.19 | 0.07 | 7.0E-11 | 0.41 |
| TG | rs6065906 | 20 | 45925376 | C | 0.19 | 0.08 | 0.04 | 0.19 | 0.05 | 2.0E-34 | 0.24 |
| TG | rs17216525 | 19 | 19551411 | T | 0.08 | -0.12 | 0.04 | 0.07 | -0.11 | 4.0E-11 | 0.47 |
| WHR | rs2371767 | 3 | 64732582 | G | 0.72 | 0.11 | 1.5E-03 | 0.73 | 0.04 | 1.6E-20 | 0.22 |
| WHR | rs1440372 | 15 | 66740813 | C | 0.72 | 0.10 | 3.5E-03 | 0.71 | 0.02 | 1.1E-10 | 0.09 |
| WHR | rs7759742 | 6 | 32413959 | A | 0.53 | 0.08 | 0.01 | 0.51 | 0.02 | 4.4E-11 | 0.10 |
| WHR | rs765751 | 1 | 219495884 | C | 0.61 | 0.08 | 0.01 | 0.61 | 0.03 | 9.0E-31 | 0.16 |
| WHR | rs8030605 | 15 | 56212400 | A | 0.12 | 0.12 | 0.01 | 0.14 | 0.03 | 8.8E-09 | 0.10 |
| WHR | rs754133 | 12 | 54025136 | A | 0.36 | 0.08 | 0.01 | 0.36 | 0.03 | 2.0E-12 | 0.16 |
| WHR | rs2071449 | 12 | 54034227 | A | 0.37 | 0.07 | 0.02 | 0.37 | 0.03 | 3.0E-14 | 0.16 |
| WHR | rs1443512 | 12 | 53948900 | A | 0.23 | 0.08 | 0.02 | 0.24 | 0.03 | 6.0E-17 | 0.13 |
| WHR | rs3936510 | 5 | 56565039 | T | 0.20 | 0.09 | 0.02 | 0.19 | 0.03 | 8.9E-09 | 0.13 |
| WHR | rs6795735 | 3 | 64719689 | C | 0.58 | 0.07 | 0.03 | 0.59 | 0.03 | 1.0E-13 | 0.17 |
| WHR | rs718314 | 12 | 26300350 | G | 0.23 | 0.08 | 0.03 | 0.26 | 0.03 | 1.0E-17 | 0.13 |
| WHR | rs10804591 | 3 | 129615390 | A | 0.79 | 0.08 | 0.03 | 0.80 | 0.03 | 6.6E-09 | 0.08 |

Abbreviations: Chromosome (Chr); base position, GRCh38 (BP); effect allele (EA); effect allele frequency (EAF).

**Supplementary Table 2**: Observed "exact" replications of SNP-phenotype associations and phenotype-specific RERs in SJLIFE and CCSS

| Cohort | Traitss | # tested / # reported SNP-phenotype associations (%) | Model[a] | Observed exact[b] replications | RER[c] (95% CI) | RER *P* |
|---|---|---|---|---|---|---|
| **SJLIFE** | Height | 519 / 535 (97.01%) | GWAS covariates only | 68 | 0.77 (0.61-0.96) | 0.02 |
| | | | +Survivor covariates | 67 | 0.76 (0.60-0.94) | 0.01 |
| | Body mass index | 123 / 126 (97.62%) | GWAS covariates only | 15 | 0.34 (0.21-0.54) | $4.9 \times 10^{-6}$ |
| | | | +Survivor covariates | 16 | 0.36 (0.23-0.57) | $1.0 \times 10^{-5}$ |
| | Waist-to-hip ratio | 73 / 74 (98.65%) | GWAS covariates only | 12 | 1.10 (0.64-1.87) | 0.73 |
| | | | +Survivor covariates | 12 | 1.10 (0.65-1.86) | 0.72 |
| | Systolic blood pressure | 67 / 74 (90.54%) | GWAS covariates only | 6 | 0.59 (0.28-1.22) | 0.15 |
| | | | +Survivor covariates | 6 | 0.59 (0.28-1.22) | 0.15 |
| | Diastolic blood pressure | 84 / 86 (97.67%) | GWAS covariates only | 3 | 0.27 (0.09-0.79) | 0.02 |
| | | | +Survivor covariates | 3 | 0.27 (0.09-0.79) | 0.02 |
| | High-density lipoprotein | 78 / 80 (97.50%) | GWAS covariates only | 18 | 0.84 (0.59-1.22) | 0.37 |
| | | | +Survivor covariates | 15 | 0.70 (0.47-1.06) | 0.09 |
| | Low-density lipoprotein | 63 / 63 (100.0%) | GWAS covariates only | 15 | 0.73 (0.51-1.06) | 0.10 |
| | | | +Survivor covariates | 15 | 0.73 (0.51-1.05) | 0.09 |
| | Total cholesterol | 73 / 73 (100.0%) | GWAS covariates only | 14 | 0.73 (0.48-1.11) | 0.14 |
| | | | +Survivor covariates | 16 | 0.84 (0.57-1.22) | 0.35 |
| | Triglycerides | 43 / 44 (97.73%) | GWAS covariates only | 13 | 1.01 (0.74-1.36) | 0.97 |
| | | | +Survivor covariates | 12 | 0.93 (0.67-1.29) | 0.66 |
| | Coronary artery disease | 92 / 94 (97.87%) | GWAS covariates only | 4 | 0.49 (0.19-1.30) | 0.15 |
| | | | +Survivor covariates | 4 | 0.49 (0.19-1.30) | 0.15 |
| | Type 2 diabetes | 98 / 102 (96.08%) | GWAS covariates only | 13 | 1.33 (0.78-2.26) | 0.30 |
| | | | +Survivor covariates | 11 | 1.12 (0.63-1.99) | 0.69 |
| | Obesity | 63 / 64 (98.44%) | GWAS covariates only | 8 | 0.38 (0.21-0.67) | $8.0 \times 10^{-4}$ |
| | | | +Survivor covariates | 8 | 0.38 (0.21-0.67) | $8.0 \times 10^{-4}$ |
| | **Total** | 1,376 / 1,415 (97.24%) | GWAS covariates only | 189 | 0.68 (0.60-0.77) | $2.4 \times 10^{-9}$ |
| | | | +Survivor covariates | 185 | 0.66 (0.58-0.76) | $4.1 \times 10^{-10}$ |
| **CCSS** | Height | 515 / 535 (96.26 %) | GWAS covariates only | 96 | 0.66 (0.55-0.77) | $6.8 \times 10^{-7}$ |
| | | | +CCS covariates | 113 | 0.77 (0.66-0.90) | $7.4 \times 10^{-4}$ |
| | Body mass index | 124 / 126 (98.41%) | GWAS covariates only | 24 | 0.43 (0.30-0.62) | $4.2 \times 10^{-6}$ |
| | | | +CCS covariates | 24 | 0.43 (0.30-0.62) | $4.2 \times 10^{-6}$ |
| | Obesity | 63 / 64 (98.44%) | GWAS covariates only | 4 | 0.15 (0.06-0.37) | $5.9 \times 10^{-5}$ |
| | | | +CCS covariates | 4 | 0.15 (0.06-0.37) | $5.9 \times 10^{-5}$ |
| | Type 2 diabetes | 96 / 102 (94.12%) | GWAS covariates only | 8 | 0.58 (0.29-1.17) | 0.13 |
| | | | +CCS covariates | 9 | 0.66 (0.34-1.26) | 0.21 |
| | Coronary artery disease | 88 / 94 (93.62%) | GWAS covariates only | 3 | 0.29 (0.09-0.90) | 0.03 |
| | | | +CCS covariates | 4 | 0.39 (0.15-1.04) | 0.06 |
| | **Total** | 886 / 921 (96.2%) | GWAS covariates only | 135 | 0.53 (0.46-0.62) | $1.1 \times 10^{-16}$ |
| | | | +CCS covariates | 154 | 0.61 (0.53-0.70) | $2.2 \times 10^{-12}$ |

Abbreviations: Number (#), Replication Enrichment Ratio (RER).

a. Results for replication analyses under two different models are reported for each phenotype: replications considering adjustment covariates typically applied in reference GWAS ("GWAS covariates only"), and replications considering both adjustment covariates typically applied in reference GWAS and adjustment covariates that should be considered in childhood cancer survivor populations ("+Survivor covariates").

b. "Exact" replication is defined as the replication of the originally reported SNP with a genome-wide significant association with the selected phenotype in reference meta-GWAS (i.e., index SNP) with $P<0.05$ and with same direction of reported association effect.

c. RER is the observed-to-expected replication ratio of genome-wide significant SNP-phenotype associations.

**Supplementary Table 3**: Expected versus observed "extended" replications of SNP-phenotype associations in SJLIFE cohort

| | | | | Extended replications (with proxy SNPs)[d] | | |
|---|---|---|---|---|---|---|
| Phenotype | SJLIFE analysis features[a] | # SNP-phenotype associations re-tested by proxy[b] | Expected replications[c] | Observed replications | RER[e] (95% CI) | RER *P* |
| Height | GWAS covariates only | 303 | 88.64 | 71 | 0.80 (0.65-0.99) | 0.04 |
| | +CCS covariates | 302 | 88.64 | 69 | 0.78 (0.63-0.97) | 0.03 |
| Body mass index | GWAS covariates only | 74 | 44.57 | 16 | 0.36 (0.23-0.57) | $1.0 \times 10^{-5}$ |
| | +CCS covariates | 75 | 44.57 | 16 | 0.36 (0.23-0.57) | $1.0 \times 10^{-5}$ |
| Waist-to-hip ratio | GWAS covariates only | 42 | 10.91 | 15 | 1.37 (0.87-2.18) | 0.18 |
| | +CCS covariates | 42 | 10.91 | 15 | 1.37 (0.87-2.17) | 0.17 |
| Systolic blood pressure | GWAS covariates only | 38 | 10.20 | 6 | 0.59 (0.28-1.22) | 0.15 |
| | +CCS covariates | 38 | 10.20 | 6 | 0.59 (0.28-1.22) | 0.15 |
| Diastolic blood pressure | GWAS covariates only | 57 | 11.12 | 3 | 0.27 (0.09-0.79) | 0.02 |
| | +CCS covariates | 57 | 11.12 | 3 | 0.27 (0.09-0.79) | 0.02 |
| High density lipoprotein | GWAS covariates only | 45 | 21.31 | 19 | 0.89 (0.62-1.28) | 0.53 |
| | +CCS covariates | 48 | 21.31 | 18 | 0.84 (0.59-1.22) | 0.37 |
| Low density lipoprotein | GWAS covariates only | 32 | 20.47 | 16 | 0.78 (0.55-1.12) | 0.18 |
| | +CCS covariates | 31 | 20.47 | 15 | 0.73 (0.51-1.05) | 0.09 |
| Total cholesterol | GWAS covariates only | 41 | 19.14 | 15 | 0.78 (0.53-1.16) | 0.22 |
| | +CCS covariates | 40 | 19.14 | 16 | 0.84 (0.57-1.22) | 0.35 |
| Triglycerides | GWAS covariates only | 25 | 12.93 | 13 | 1.01 (0.74-1.36) | 0.97 |
| | +CCS covariates | 26 | 12.93 | 12 | 0.93 (0.67-1.29) | 0.66 |
| Coronary artery disease | GWAS covariates only | 52 | 8.16 | 6 | 0.74 (0.33-1.63) | 0.45 |
| | +CCS covariates | 53 | 8.16 | 6 | 0.74 (0.33-1.63) | 0.45 |
| Type 2 diabetes | GWAS covariates only | 65 | 9.80 | 13 | 1.33 (0.78-2.26) | 0.30 |
| | +CCS covariates | 67 | 9.80 | 12 | 1.23 (0.71-2.12) | 0.47 |
| Obesity | GWAS covariates only | 38 | 21.25 | 8 | 0.38 (0.21-0.67) | $8.1 \times 10^{-4}$ |
| | +CCS covariates | 38 | 21.25 | 8 | 0.38 (0.21-0.67) | $8.1 \times 10^{-4}$ |
| All SNP-phenotype combinations | GWAS covariates only | 812 | 278.49 | 201 | 0.72 (0.64-0.82) | $2.2 \times 10^{-7}$ |
| | +CCS covariates | 817 | 278.49 | 196 | 0.70 (0.62-0.80) | $3.2 \times 10^{-8}$ |

Abbreviations: Number (#), childhood cancer survivor (CCS), Replication Enrichment Ratio (RER).

a. Results for two different replication analyses in SJLIFE are reported: replications considering adjustment covariates typically applied in reference GWAS ("GWAS covariates only"), and replications considering both adjustment covariates typically applied in reference GWAS and adjustment covariates that should be considered in childhood cancer survivor populations ("+CCS covariates").

b. Number of SNP associations that were not replicated with index SNPs and re-tested using best SNP proxies for index SNPs, when available; best proxies were defined as the SNPs in high LD ($r^2$>0.8) in the 1000 Genomes European reference population likely to fall in the same LD block (within a 5-kb window of the index SNP).

c. Expected numbers of replications are estimated by power calculations and the observed SNP effect allele frequency; power calculations assume an additive genetic inheritance model, α=0.05, and effect sizes based on reference meta-GWAS.

d. "Extended" replication is defined as the replication of either the index SNP or a best SNP proxy with *P*<0.05 and with same direction of reported association effect.

e. RER is the observed-to-expected replication ratio of genome-wide significant SNP-phenotype associations in the SJLIFE cohort.

**Supplementary Table 4:** Expected versus observed "exact" replications of independent SNP-phenotype associations in SJLIFE cohort

| | | | Exact replications[d] | | | |
|---|---|---|---|---|---|---|
| Phenotype | # independent SNP-phenotype associations[a] | Expected replications[b] | SJLIFE analysis features[c] | Observed replications | RER[e] (95% CI) | RER *P* |
| Height | 431 | 65.72 | GWAS covariates only | 51 | 0.78 (0.60-1.00) | 0.05 |
| | | | +CCS covariates | 51 | 0.78 (0.60-1.00) | 0.05 |
| Body mass index | 100 | 29.56 | GWAS covariates only | 9 | 0.30 (0.16-0.57) | $2.3 \times 10^{-4}$ |
| | | | +CCS covariates | 10 | 0.34 (0.18-0.62) | $4.4 \times 10^{-4}$ |
| Waist-to-hip ratio | 60 | 8.48 | GWAS covariates only | 11 | 1.30 (0.75-2.25) | 0.35 |
| | | | +CCS covariates | 11 | 1.30 (0.76-2.22) | 0.34 |
| Systolic blood pressure | 64 | 9.65 | GWAS covariates only | 5 | 0.52 (0.23-1.17) | 0.11 |
| | | | +CCS covariates | 5 | 0.52 (0.23-1.17) | 0.11 |
| Diastolic blood pressure | 77 | 9.72 | GWAS covariates only | 2 | 0.21 (0.05-0.79) | 0.02 |
| | | | +CCS covariates | 2 | 0.21 (0.05-0.79) | 0.02 |
| High-density lipoprotein | 72 | 17.75 | GWAS covariates only | 15 | 0.84 (0.57-1.26) | 0.41 |
| | | | +CCS covariates | 12 | 0.68 (0.43-1.07) | 0.10 |
| Low-density lipoprotein | 58 | 17.37 | GWAS covariates only | 12 | 0.69 (0.46-1.03) | 0.07 |
| | | | +CCS covariates | 12 | 0.69 (0.47-1.02) | 0.07 |
| Total cholesterol | 73 | 19.14 | GWAS covariates only | 14 | 0.73 (0.48-1.11) | 0.14 |
| | | | +CCS covariates | 16 | 0.84 (0.57-1.22) | 0.35 |
| Triglycerides | 40 | 11.32 | GWAS covariates only | 11 | 0.97 (0.70-1.35) | 0.86 |
| | | | +CCS covariates | 10 | 0.88 (0.62-1.27) | 0.50 |
| Coronary artery disease | 67 | 5.62 | GWAS covariates only | 2 | 0.36 (0.09-1.41) | 0.14 |
| | | | +CCS covariates | 2 | 0.36 (0.09-1.40) | 0.14 |
| Type 2 diabetes | 95 | 9.46 | GWAS covariates only | 12 | 1.27 (0.73-2.21) | 0.40 |
| | | | +CCS covariates | 10 | 1.06 (0.58-1.93) | 0.86 |
| Obesity | 48 | 14.44 | GWAS covariates only | 3 | 0.21 (0.07-0.59) | $3.1 \times 10^{-3}$ |
| | | | +CCS covariates | 3 | 0.21 (0.07-0.59) | $3.1 \times 10^{-3}$ |
| All SNP-phenotype associations | 1185 | 218.23 | GWAS covariates only | 147 | 0.67 (0.58-0.78) | $1.0 \times 10^{-7}$ |
| | | | +CCS covariates | 144 | 0.66 (0.57-0.76) | $2.9 \times 10^{-8}$ |

Abbreviations: Number (#), childhood cancer survivor (CCS), Replication Enrichment Ratio (RER).

a. For each phenotype, the compiled set of SNP-phenotype associations was "pruned" to the set of independent SNPs (not in high LD [$R^2 \geq 0.8$] with any SNP within a 500-kb window in the 1000G European reference population); the SNP with the highest effect allele frequency in SJLIFE was the retained SNP among each cluster of high-LD SNPs.

b. Expected numbers of replications are estimated by power calculations and the observed SNP effect allele frequency; power calculations assume an additive genetic inheritance model, $\alpha = 0.05$, and effect sizes based on reference meta-GWAS.

c. Results for two different replication analyses in SJLIFE are reported: replications considering adjustment covariates typically applied in reference GWAS ("GWAS covariates only"), and replications considering both adjustment covariates typically applied in reference GWAS and adjustment covariates that should be considered in childhood cancer survivor populations ("+CCS covariates").

d. "Exact" replication is defined as the replication of the originally reported SNP with a genome-wide significant association with the selected phenotype in reference meta-GWAS (i.e., index SNP) with $P<0.05$ and with same direction of reported association effect.

e. RER is the observed-to-expected replication ratio of genome-wide significant SNP-phenotype associations in the SJLIFE cohort.

**Supplementary Table 5:** Expected versus observed replications of SNP-phenotype associations using a Bonferroni-corrected P-value threshold to define replication in the SJLIFE cohort

| Phenotype | Sample size (N) | Bonferroni-corrected significance | Expected replications[a] | SJLIFE analysis features[b] | Observed replications (P<threshold) |
|---|---|---|---|---|---|
| Height | 2,025 | $\alpha=9.6x10^{-5}$ | 2.14 | GWAS covariates only<br>+CCS covariates | 0<br>0 |
| Body mass index | 2,229 | $\alpha=4.1x10^{-4}$ | 20.50 | GWAS covariates only<br>+CCS covariates | 7<br>4 |
| Waist-to-hip ratio | 2,204 | $\alpha=6.8x10^{-4}$ | 0.54 | GWAS covariates only<br>+CCS covariates | 0<br>0 |
| Systolic blood pressure | 2,020 | $\alpha=7.5x10^{-4}$ | 0.59 | GWAS covariates only<br>+CCS covariates | 0<br>0 |
| Diastolic blood pressure | 2,020 | $\alpha=6.0x10^{-4}$ | 0.46 | GWAS covariates only<br>+CCS covariates | 0<br>0 |
| High density lipoprotein | 1,984 | $\alpha=6.4x10^{-4}$ | 6.31 | GWAS covariates only<br>+CCS covariates | 3<br>3 |
| Low density lipoprotein | 1,964 | $\alpha=7.9x10^{-4}$ | 7.57 | GWAS covariates only<br>+CCS covariates | 5<br>5 |
| Total cholesterol | 1,997 | $\alpha=6.8x10^{-4}$ | 4.86 | GWAS covariates only<br>+CCS covariates | 5<br>5 |
| Triglycerides | 1,997 | $\alpha=1.2x10^{-3}$ | 4.93 | GWAS covariates only<br>+CCS covariates | 3<br>3 |
| Coronary artery disease | 2,079 | $\alpha=5.4x10^{-4}$ | 0.24 | GWAS covariates only<br>+CCS covariates | 0<br>0 |
| Type 2 diabetes | 2,112 | $\alpha=5.1x10^{-4}$ | 0.58 | GWAS covariates only<br>+CCS covariates | 0<br>0 |
| Obesity | 2,229 | $\alpha=7.9x10^{-4}$ | 5.60 | GWAS covariates only<br>+CCS covariates | 2<br>1 |

Abbreviations: Number (#), childhood cancer survivor (CCS), Replication Enrichment Ratio (RER).

a. Expected numbers of replications are estimated by power calculations and the observed SNP effect allele frequency; power calculations assume an additive genetic inheritance model, α=Bonferroni-corrected significance level, and effect sizes based on reference meta-GWAS.

b. Results for two different replication analyses in SJLIFE are reported: replications considering adjustment covariates typically applied in reference GWAS (“GWAS covariates only”), and replications considering both adjustment covariates typically applied in reference GWAS and adjustment covariates that should be considered in childhood cancer survivor populations (“+CCS covariates”).

**Supplementary Table 6**: Top cell/tissue types with significant (FDR≤0.05) GTEx *cis*-eQTL enrichments among SNPs with SNP-phenotype association replications (N=170) compared to SNPs without SNP-phenotype association replications (N=1,061) in SJLIFE

| Tissue | # significant eQTLs, replicated SNPs (total: 1,659 significant eQTLs) | # significant eQTLs, non-replicated SNPs (total: 14,960 significant eQTLs) | Enrichment OR | P |
|---|---|---|---|---|
| Liver | 22 | 121 | 1.65 | 0.04 |
| Adipose, Visceral Omentum | 61 | 413 | 1.34 | 0.04 |
| Adipose, Subcutaneous | 100 | 690 | 1.33 | 0.01 |
| Testis | 74 | 520 | 1.30 | 0.04 |

**Supplementary Table 7**: Top cell/tissue types with 15-state ChromHMM enhancer (*P*<0.05) and promoter (*P*<0.1) overlap enrichments among replicated SNPs (N=170 SNPs) compared to non-replicated SNPs (N=1,061 SNPs) in SJLIFE

| Chromatin State | Epigenome Name (EID) | # chromatin state overlaps, replicated SNPs (N=170) | # chromatin state overlaps, non-replicated SNPs (N=1061) | Enrichment OR | P |
|---|---|---|---|---|---|
| Enhancer | Fetal kidney (E086) | 18 | 47 | 2.55 | $2.4 \times 10^{-3}$ |
| Enhancer | Brain cingulate gyrus (E069) | 22 | 69 | 2.14 | 0.01 |
| Enhancer | Sigmoid colon (E106) | 17 | 55 | 2.03 | 0.02 |
| Enhancer | Brain angular gyrus (E067) | 18 | 60 | 1.97 | 0.03 |
| Enhancer | Brain dorsolateral prefrontal cortex (E073) | 16 | 54 | 1.94 | 0.03 |
| Enhancer | Brain inferior temporal lobe | 17 | 58 | 1.92 | 0.04 |
| Enhancer | H1 BMP4 derived trophoblast | 24 | 86 | 1.86 | 0.02 |
| Enhancer | Brain hippocampus middle (E071) | 23 | 84 | 1.82 | 0.03 |
| Enhancer | Adipose nuclei (E063) | 28 | 112 | 1.67 | 0.04 |
| Enhancer | H1 derived mesenchymal stem cells (E006) | 25 | 100 | 1.66 | 0.04 |
| Enhancer | Breast myoepithelial (E027) | 32 | 132 | 1.63 | 0.03 |
| Promoter | Mesenchymal stem cell derived adipocyte (E023) | 13 | 48 | 1.75 | 0.09 |
| Promoter | HepG2 hepatocellular carcinoma (E118) | 13 | 48 | 1.75 | 0.09 |
| Promoter | Osteoblast (E129) | 13 | 48 | 1.75 | 0.09 |

Color coding:
Green: anthropometric phenotypes
Blue: blood pressure phenotypes
Yellow: blood lipid phenotypes
Pink: cardiovascular/metabolic disease

**Supplementary Table 8**: Significant (FDR<0.05) *cis*-meQTLs from BIOS QTL for 1,231 investigated meta-GWAS SNPs

| | % (N) |
|---|---|
| **Significant *cis*-meQTLs in BIOS QTL (≤250 kb SNP-CpG distance)** | |
| Meta-GWAS SNPs with ≥1 significant *cis*-meQTLs | 87.5% (1,077) |
| Non-replicated SNPs with ≥1 significant *cis*-meQTL (N=1,061) | 87.5% (928) |
| Replicated SNPs with ≥1 significant *cis*-meQTL (N=170) | 87.6% (149) |
| Significant *cis*-meQTLs for non-replicated SNPs (N=19,091) | 86.4% (16,486) |
| Significant *cis*-meQTLs for replicated SNPs (N=19,091) | 13.6% (2,605) |

Abbreviations: Methylation quantitative trait loci (meQTL).

**Supplementary Table 9:** Enrichment of SJLIFE-validated[a] *cis*-meQTLs (N=236)

| SNP-replication groupings and meQTL types | # SNPs with SJLIFE-validated *cis*-meQTLs | # SNPs with established[d] *cis*-meQTLs | % | # SNPs with SJLIFE-validated *cis*-meQTLs | # SNPs with established[d] *cis*-meQTLs | % | OR | 95% CI | Enrichment or depletion *P* |
|---|---|---|---|---|---|---|---|---|---|
| Non-replicated vs. replicated[b] | Non-replicated SNPs | | | Replicated SNPs | | | | | |
| *cis*-meQTLs validated in SJLIFE, P<0.05 | 783 | 928 | 84.4% | 114 | 149 | 76.5% | 1.66 | 1.06-2.55 | 0.02 |
| Treatment-sensitive vs treatment-insensitive[c] | Treatment-sensitive SNPs | | | Treatment-insensitive SNPs | | | | | |
| *cis*-meQTLs validated in SJLIFE, P<0.05 | 38 | 42 | 90.5% | 37 | 57 | 64.9% | 5.06 | 1.50-22.30 | $4.1 \times 10^{-3}$ |

Abbreviations: Number (#), methylation quantitative trait loci (meQTL)

a. Validated *cis*-meQTLs were defined as *cis*-meQTLs nominated in BIOS QTL (FDR<0.05) that showed SNP-CpG methylation associations with *P*<0.05 in SJLIFE and had the same direction of allelic effect as BIOS QTL.
b. Non-replicated SNPs have no replicated SNP-phenotype associations in our main SJLIFE analysis, while replicated SNPs have at least one replicated association.
c. Treatment-insensitive SNPs have SNP-phenotype association replications in treatment-unexposed and treatment-exposed samples in SJLIFE, while treatment-sensitive SNPs do not have SNP-phenotype association replications in our main analysis but showed replicated associations in samples not exposed to treatments in SJLIFE.
d. “Established” *cis*-meQTLs are significant *cis*-meQTLs (FDR<0.05) in BIOS QTL.

**Supplementary Table 10:** Treatment exposures in experimental SJLIFE sample with methylation and genotype data (N=236)

| Cancer treatment exposures | Total | % Exposed[a] (N) |
|---|---|---|
| Cranial radiotherapy | 229 | 40.2% (92) |
| Chest radiotherapy | 229 | 34.5% (79) |
| Abdominal radiotherapy | 229 | 33.2% (76) |
| Pelvic radiotherapy | 229 | 27.9% (64) |
| Anthracyclines | 235 | 52.3% (123) |
| Corticosteroids | 234 | 42.3% (99) |
| Carboplatin | 236 | 3.4% (8) |
| Cisplatin | 236 | 8.9% (21) |

a. Exposed to radiotherapy was defined as scatter dose >20 cGy; exposed to chemotherapy was defined as dose >0.

**Supplementary Table 11:** Discordant SNP-methylation and treatment-methylation associations at CpGs for validated *cis*-meQTLs in SJLIFE

| | Non-replicated CpGs[a] (3,604 CpGs) | | | Replicated CpGs[b] (549 CpGs) | | | | | |
|---|---|---|---|---|---|---|---|---|---|
| Treatment definition | # CpGs with treatment association[c] (P<0.05) | # Discordant[d] | % | # CpGs with treatment association[c] (P<0.05) | # Discordant[d] | % | OR | 95% CI | Enrichment or depletion P |
| Radiotherapies | | | | | | | | | |
| Cranial RT | 176 | 90 | 51.1% | 27 | 17 | 63.0% | 0.62 | 0.24-1.52 | 0.30 |
| Chest RT | 315 | 151 | 47.9% | 71 | 18 | 25.4% | 2.70 | 1.48-5.14 | $5.3x10^{-4}$ |
| Abdominal RT | 240 | 132 | 55.0% | 41 | 16 | 39.0% | 1.91 | 0.92-4.03 | 0.06 |
| Pelvic RT | 294 | 159 | 54.1% | 52 | 15 | 28.8% | 2.90 | 1.48-5.94 | $8.7x10^{-4}$ |
| Chemotherapies | | | | | | | | | |
| Anthracyclines | 203 | 94 | 46.3% | 28 | 17 | 60.7% | 0.56 | 0.22-1.34 | 0.16 |
| Corticosteroids | 188 | 95 | 50.5% | 34 | 20 | 58.8% | 0.72 | 0.31-1.59 | 0.46 |
| Cisplatin | 220 | 101 | 45.9% | 29 | 17 | 58.6% | 0.60 | 0.25-1.41 | 0.24 |

Abbreviations: Radiotherapy (RT); number (#).

a. Non-replicated CpGs are CpGs linked to SJLIFE-validated *cis*-meQTLs for non-replicated SNPs.
b. Replicated CpGs are CpGs linked to SJLIFE-validated *cis*-meQTLs for replicated SNPs.
c. CpGs with treatment associations are CpGs linked to SJLIFE-validated *cis*-meQTLs that are also associated with treatment dose for the specified treatmet ($P$<0.05).
d. Discordant CpGs have directionally discordant (conflicting) SNP-methylation and treatment-methylation associations.

**Supplementary Table 12:** Reported associations between rs1552224 (chr11:72722053, GRCh38 build) and T2D risk and replication results in subgroups of SJLIFE survivors stratified by exposures to abdominal or pelvic radiation therapy

| Study sample | Sample size, EUR | % T2D cases ($N_{cases}$) | EA | EAF | Beta | SE | *P*-value |
|---|---|---|---|---|---|---|---|
| Reference GWAS: Zhao *et al.*, 2017 | 20,298 | 19.1% (3,871) | A | 0.85 | 0.10 | 0.01 | $1.4x10^{-13}$ |
| Reference GWAS: Voight *et al.*, 2010 | 141,454 | 30.1% (42,542) | A | 0.88 | 0.13 | 0.01 | $1.4x10^{-22}$ |
| SJLIFE: All survivors | 2,112 | 7.1% (149) | A | 0.84 | 0.24 | 0.19 | 0.21 |
| SJLIFE: Treatment subgroup | 1,164 | 9.0% (105) | A | 0.84 | -0.04 | 0.21 | 0.85 |
| SJLIFE: No-treatment subgroup | 948 | 4.6% (44) | A | 0.85 | 1.74 | 0.63 | $5.9x10^{-3}$ |

Abbreviations: EUR (European ancestry); % (percent); T2D (type 2 diabetes); EA (effect allele); EAF (effect allele frequency); SE (standard error).
Beta values are given as ln(odds ratio). “Treatment” refers to any exposure to abdominal or pelvic radiation therapy.
